# Evolving cryo-EM structural approaches for GPCR drug discovery

**DOI:** 10.1101/2021.01.11.426276

**Authors:** Xin Zhang, Rachel M. Johnson, Ieva Drulyte, Lingbo Yu, Abhay Kotecha, Radostin Danev, Denise Wootten, Patrick M. Sexton, Matthew J. Belousoff

**Author notes:** Correspondence to: Denise Wootten, Patrick M. Sexton, Matthew J. Belousoff.

## Abstract

G protein-coupled receptors (GPCRs) are the largest class of cell surface drug targets. Advances in biochemical approaches for the stabilisation of GPCR:transducer complexes together with improvements in the technology and application of cryo-EM has recently opened up new possibilities for structure-assisted drug design of GPCR agonists. Nonetheless, limitations in the commercial application of some of these approaches, including the use of nanobody 35 (Nb35) for stabilisation of GPCR:Gs complexes, and the high cost of 300kV imaging have restricted broad application of cryo-EM in drug discovery. Here, using the PF 06882961-bound GLP-1R as exemplar, we validated formation of stable complexes with a modified Gs protein in the absence of Nb35 that had equivalent resolution in the drug binding pocket to complexes solved in the presence of Nb35, while the G protein displayed increased conformational dynamics. In parallel, we assessed the performance of 200kV versus 300kV image acquisition using a Falcon 4 or K3 direct electron detector. We show that with 300kV Krios, both bottom mounted Falcon 4 and energy filtered (25eV slit) Bio-Quantum K3 produced similar resolution. Moreover, the 200kV Glacios with bottom mounted Falcon 4 yielded a 3.2 Å map with clear density for bound drug and multiple structurally ordered waters. Our work paves the way for broader commercial application of cryo-EM for GPCR drug discovery.

## Introduction

Advances in single particle cryo-electron microscopy (cryo-EM) have enabled high resolution structure determination for previously intractable proteins and protein complexes. Among these are integral membrane proteins including receptor and ion channels that are key drug target classes, with G protein-coupled receptors (GPCRs) the largest superfamily of cell surface receptor proteins^1,2^.

While X-ray crystallography has been important in GPCR structure elucidation, particularly for inhibitor-bound structures, cryo-EM has heralded a new era in determination of agonist-bound GPCRs in the fully active state where they are coupled to canonical transducer proteins. Since the first cryo-EM GPCR structure, reported in 2017 at moderate resolution (~4 Å)^3^, continuing advances in the biochemical tools for complex stabilisation, detector technology, image acquisition and cryo-EM data processing have enabled progressive improvements in resolution of structures and the spectrum of GPCRs and transducers that can solved^4^. Nonetheless, while resolutions that can support accurate modelling of drug-receptor interactions have become more common, there remain substantial barriers to routine use of cryo-EM for commercial GPCR drug discovery and development. Among these are technology patents around tools for stabilisation of GPCR-G protein complexes and the high cost of access for state-of-the-art 300kV imaging.

The primary functional role of GPCRs is catalysation of guanine nucleotide exchange by transducer G proteins following agonist promoted recruitment to the receptor, leading to activation of the G protein and dissociation^5^. Consequently, the interaction between receptor and G protein is inherently unstable and stabilisation of the ternary complex of agonist-GPCR-G protein is required for structure determination. One of the most important signal transducers for GPCRs is the Gs protein. A key tool for stabilisation of GPCR-Gs complexes has been nanobody 35 (Nb35) that binds across the interface of Gαs and Gβ subunits, which was first applied to the β_2_-adrenergic receptor-Gs complex and enabled the first ternary GPCR complex structure to be determined^6^. Alternate or supplementary technologies have subsequently been introduced, including mutant Gα proteins^7^, nanoBit tethering^8,9^ and mini-Gα proteins that reduce or eliminate guanine nucleotide binding and/or stabilise the interaction between heterotrimeric subunits^10–12^. In practice, these have primarily been used in combination with Nb35 for Gs protein complexes to provide enhanced complex stability. Both Nb35 and mini-G proteins are protected technologies (WO2012175643A2; WO2017129998) that require licensing for commercial application, while the utility of mutant (commonly referred to as dominant negative (DN)) Gαs protein alone for stabilisation of ternary complexes and structure determination has not been explored.

The glucagon-like peptide-1 receptor (GLP-1R) is a major, validated target for treatment of type 2 diabetes and obesity, with numerous approved peptide therapeutics^13^. The emergence of small molecule GLP-1R agonists that can be taken orally has sparked renewed interest in discovery and development of novel GLP-1R drugs. Among the most promising is PF 06882961, and we recently reported the structure of this compound bound to the active GLP-1R that had high-resolution in the binding pocket, including numerous structural waters^14^. As with most Gs protein-coupled complexes, Nb35 was used as part of the stabilisation strategy. In the current study we have used PF 06882961-GLP-1R to examine the stability and structural resolution of complexes formed in the absence of Nb35, using DNGαs as the stand-alone stabilising technology. Moreover, we have compared structural resolution of this complex following imaging with 200kV or 300kV cryo-EM supported by the latest detector technology (Gatan K3 or Thermo Fisher Scientific Falcon 4). We reveal that while dynamics of the G protein are increased in the absence of Nb35, the resolution of the small molecule binding pocket is similar to that observed when Nb35 was present. Of note, the resolution of the compound binding pocket in complexes imaged with a 200kV Glacios-Falcon 4 was qualitatively similar to that achieved with either Krios-K3 or Krios-Falcon 4 imaged complexes.

## Results

### Purification and structure determination of PF 06882961-GLP-1R-DNGs

Similar to previously described complexes of PF 06882961-GLP-1R-DNGs-Nb35^14^, ternary complexes formed in the absence of Nb35 demonstrated robust stability that could be sequentially purified to homogeneity by affinity chromatography followed by SEC (**Suppl. Fig. 1A**), with each of the components of the complex verified by Coomassie stained SDS-PAGE and western blot analysis probing for His tags (GLP-1R, Gβ1) or Gαs (**Suppl. Fig. 1B**), and stability of the complex by negative stain TEM (**Suppl. Fig. 1C, 1D**). Vitrified samples of PF 06882961-GLP-1R-DNGs complex were prepared and imaged by cryo-TEM on a Krios G3 300kV microscope, with data collected on a K3 detector (**Suppl. Fig. 1E**). 2D classification revealed well-resolved secondary structure features and different projections of the particles (**Suppl. Fig. 1F**), with a consensus 3D reconstruction of 2.9 Å resolution at gold standard FSC 0.143 (**Fig. 1A, 1B; Suppl. Fig. 1G**) from ~386K particles. While there was substantial preferred orientation of the sample, there was broad angular coverage at lower frequency (**Suppl. Fig. 1H**). The alpha helical domain (AHD) of Gαs subunit, and parts of Gβ1 and Gγ2, were poorly resolved with limited density in the consensus map (**Fig. 1A, left panels**). The transmembrane domain (TM) of the receptor and G protein core have relatively higher resolution < 3.0 Å (**Fig. 1A**), allowing clear rotamer placements for most amino acids within these regions (**Suppl. Fig. 2**). To improve the resolution of domains with higher mobility, local 3D refinements focused on the receptor and G protein were performed in RELION, yielding clearer densities with local resolution ranging from 2.9 – 4.1 Å (**Fig. 1A, right panels**). Most regions with higher mobility were well resolved in these local refined maps, facilitating robust molecular modelling, including the receptor ECD, ECD/TM1 linker, loop regions (extracellular and intracellular loops, ECLs and ICLs respectively), ligand binding site, αN helix of Gα, C-terminal helix of Gβ and Gγ subunit (**Fig. 1A–1C; Suppl. Fig. 2**). Nonetheless, there was limited density of the receptor ICL3 (residues L339^ICL3^ – K342^ICL3^) and residues within the β3/α2 loop of the Gαs subunit (A226^β3/α2_loop^ – E330^β3/α2_loop^), allowing only backbone level modelling for these regions. The density of residues N115^ECD^ – S117^ECD^ in the receptor ECD was discontinuous and deleted from the final model (**Fig. 1C**).

**Figure 1.**
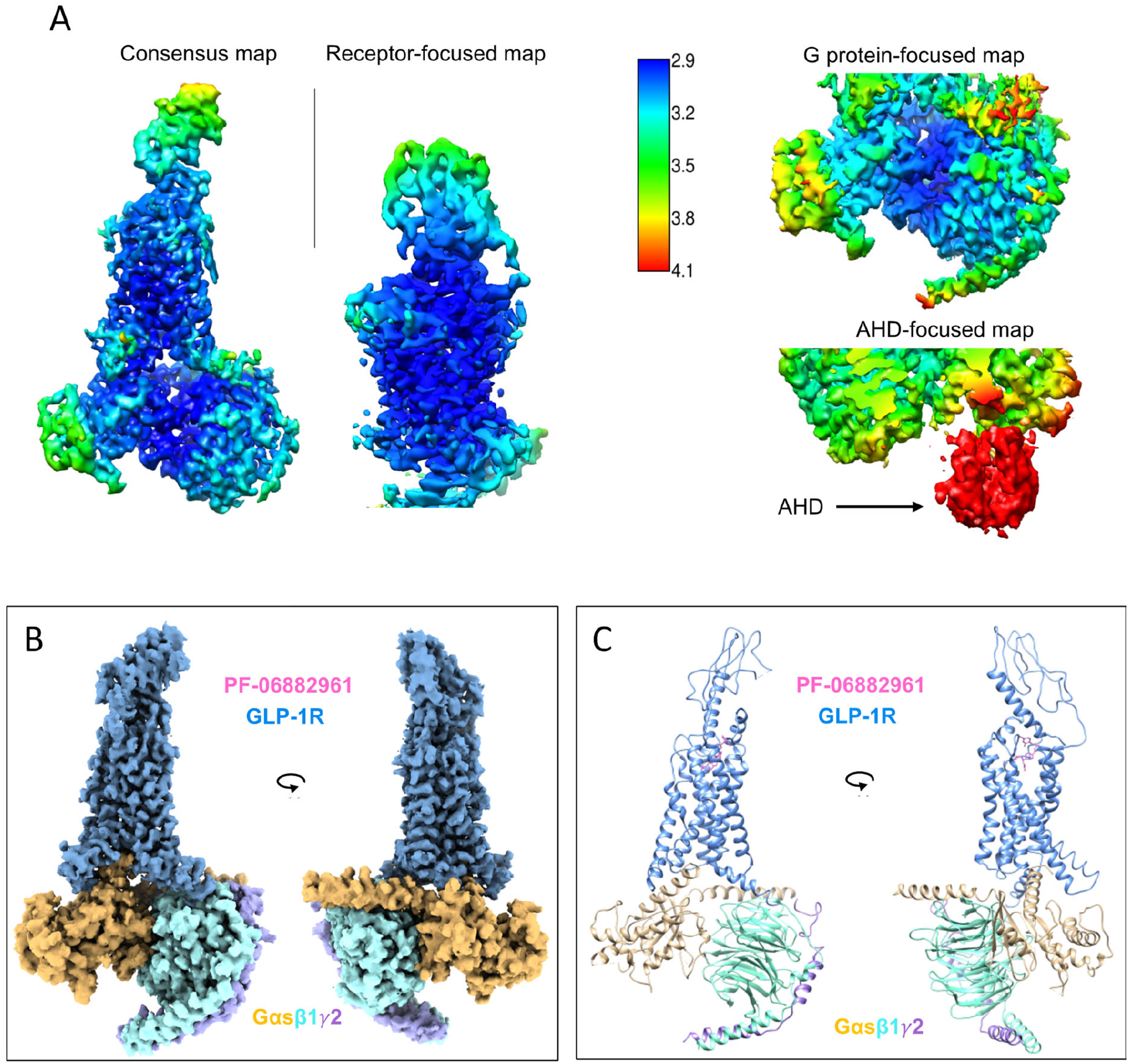
Cryo-EM structure of PF 06882961-GLP-1R-DNGs complex. (**A**) Local resolution-filtered EM maps (consensus and receptor/G protein focused refinements) displaying local resolution (Å) coloured from highest resolution (dark blue) to lowest resolution (red). (**B**) Orthogonal view of the cryo-EM map generated from consensus and local refined maps via zone of 2 Å and mask on each component of complex modelled into the maps using UCSF ChimeraX; (**C**) Backbone models of PF 06882961-GLP-1R-Gs complex in ribbon format. Colouring denotes different segments as highlighted on the figure panels.

### Comparison of PF 06882961 binding to GLP-1R-DNGs complexes in the presence and absence of Nb35

The PF 06882961 compound in the GLP-1R-DNGs complex adopted an equivalent binding pose with identical interactions with the surrounding receptor residues to that previously reported in the complex formed in the presence of Nb35 (PDB: 6X1A)^14^ (**Fig. 2A, 2B; Suppl. Fig. 3; Table S1**). Although there were minor differences in the location of water molecules below the binding pocket in the consensus map, the overlap in rotamer orientation of residues within the water-mediated polar network and the central polar core (**Fig. 2**) indicated that the presence or absence of Nb35 during structure determination did not alter either the active receptor conformation or the binding mode for complexes with PF 06882961 (**Suppl. Fig. 4A, 4B**). Similarly, there was also a high degree of overlap in the conformation of the intracellular face of the receptor, with only minor differences observed in the conformation of ICL2 and the C terminal end of H8, as well as an outward shift of ~2 Å of the intracellular tip of TM6 when measured at the Cα of T343^6.32^ relative to the Nb35-stabilised complex; this is accompanied by a small outward movement of ICL3 (**Suppl. Fig. 4**).

**Figure 2.**
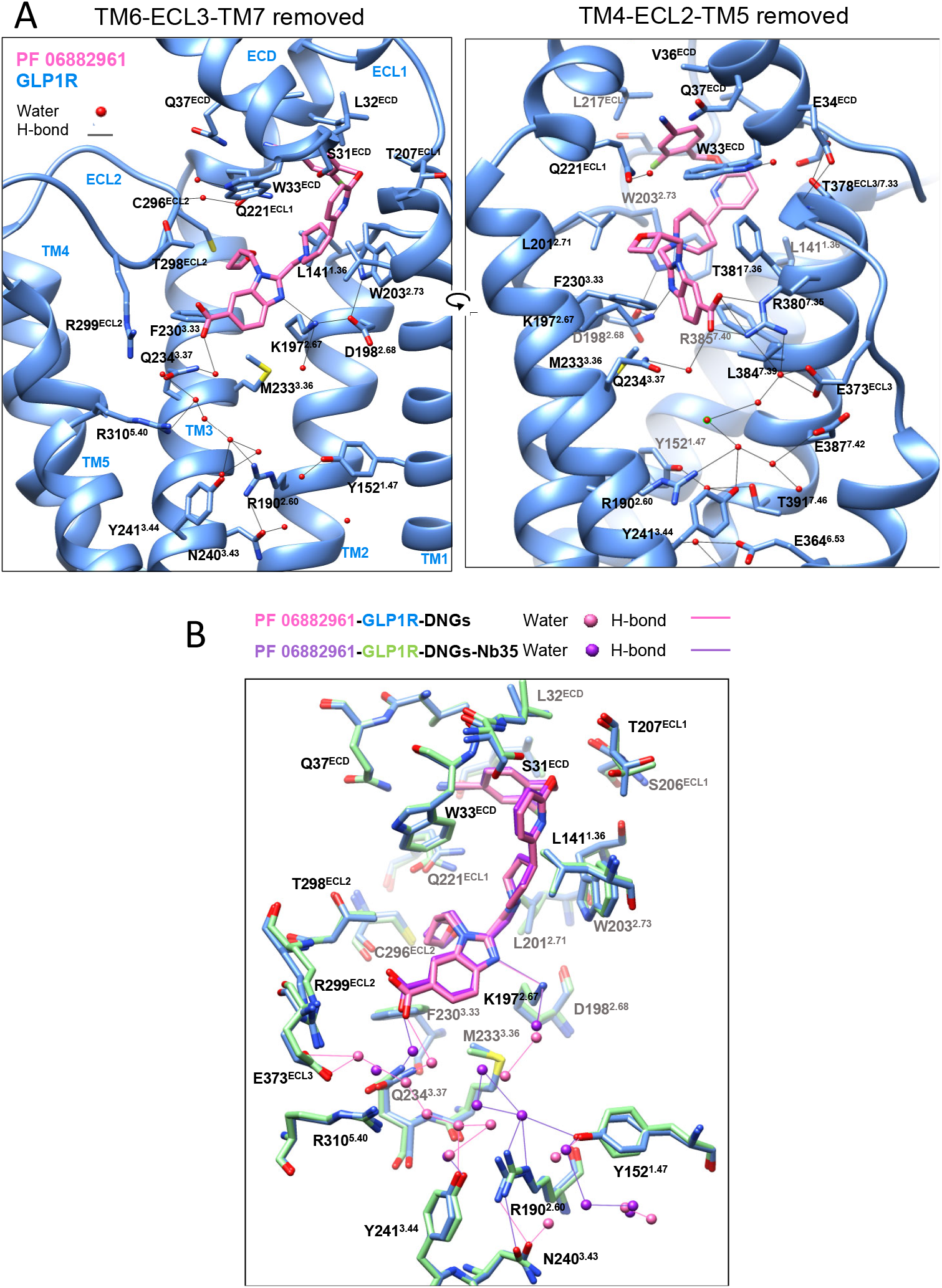
Interactions of PF 06882961 within the binding cavity of the GLP-1R. (**A**) Side view of PF 06882961 within the binding site of the GLP-1R TM bundle. Left: viewed from the upper portion of TM6/TM7 where TM6-ECL3-TM7 have been removed for clarity; Right: viewed from the upper portion of TM4/TM5 where TM4-ECL2-TM5 have been removed for clarity. Black lines depict hydrogen bonds as determined using UCSF Chimera. GLP-1R residues that interact with PF 06882961 (stick representation) or waters (red spheres) within the GLP-1R binding cavity are displayed in stick format coloured by heteroatom, with the backbone of the receptor in ribbon format; Colouring denotes different components as highlighted on the figure panels. (**B**) Comparison of the PF 06882961 binding mode in complexes formed in the presence or absence of Nb35. Overlay of the PF 06882961 binding site of the complex in the presence (PDB: 6X1A) or absence of Nb35. GLP-1R residues that interact with PF 06882961 or waters (spheres) within the GLP-1R binding cavity are displayed in stick format coloured by heteroatom. Colouring denotes different components as highlighted on the figure panel.

### Gs protein interactions with GLP-1R in the presence and absence of Nb35

Not surprisingly, complexes formed in the absence of Nb35 had a very similar pattern of interaction between the GLP-1R and Gs to that observed for complexes with Nb35 (**Fig. 3A–3C**; **Table S2**)^14^, forming primarily hydrophobic interactions with Gαs, including between the intracellular part of TM2/3/5/6 and ICL2/3 of the receptor and the α5 helix and αN-β1 junction of Gαs (**Fig. 3A, 3B; Suppl. Fig. 5A, 5B**). As noted above, there were differences in the conformation of ICL2 leading to alterations to the observed interactions with Gαs (**Fig. 3B; Table S2**). In the consensus map without Nb35, F257^ICL2/3.60^ located within the junction of αN and α5 helices, interacted extensively with V217^β3^, F376^α5H^ and R380^α5H^ of the α5 helix and H41^αN^ of the αN helix via van der Waals, whereas interactions are reduced for V217^β3^, F376^α5H^ and H41^αN^ in the Nb35-stabilised structure (**Suppl. Fig. 5A, 5B; Table S2**). Interestingly, there was an additional density present in the EM map at lower contour that could support alternative modelling of ICL2, in which F257^ICL2/3.60^ is oriented away from the αN-α5 junction (**Suppl. Fig. 5C, left panel**). The potential for alternate conformations of ICL2 was also supported by additional density in the Nb35-stabilized structure, although the density for ICL2 was less robust than that in the complex without Nb35 (**Suppl. Fig. 5C, right panel**). Direct hydrogen bonds occurred between the GLP-1R and Gαs that included Q384^α5H^ with L255^3.58^/K334^5.64^, R385^α5H^ with K334^5.64^/N338^ICL3^ and E392^α5H^ with R348^6.37^, while there was a water-mediated interaction between Y391^α5H^ and E247^3.50^ (**Suppl. Fig. 5B**). While most of these interactions were conserved in the Nb35-stabilized PF 06882961 and GLP-1 complexes, the shift in the position of TM6 noted above caused small changes to the receptor-Gαs interface (**Fig. 3A–3C; Table S2**). In contrast, the GLP-1R made fewer interactions with the Gβ1 in the absence of Nb35 than in the Nb35-stabilized structure where only the π-π stacking between R52^β^ of Gβ1 and R170^ICL1^ was conserved. Although the backbone of H8 aligned well between two structures, the residues K415^8.56^ and R419^8.60^ had distinct side chain rotamer placements that combined with a small translational shift in the position of the Gβ subunit led to loss of interaction of H8 residues with A309^β^, G310^β^, H311^β^ and D312^β^ that are observed in the Nb35-stabilised consensus structure (**Fig. 3C; Table S2**).

**Figure 3.**
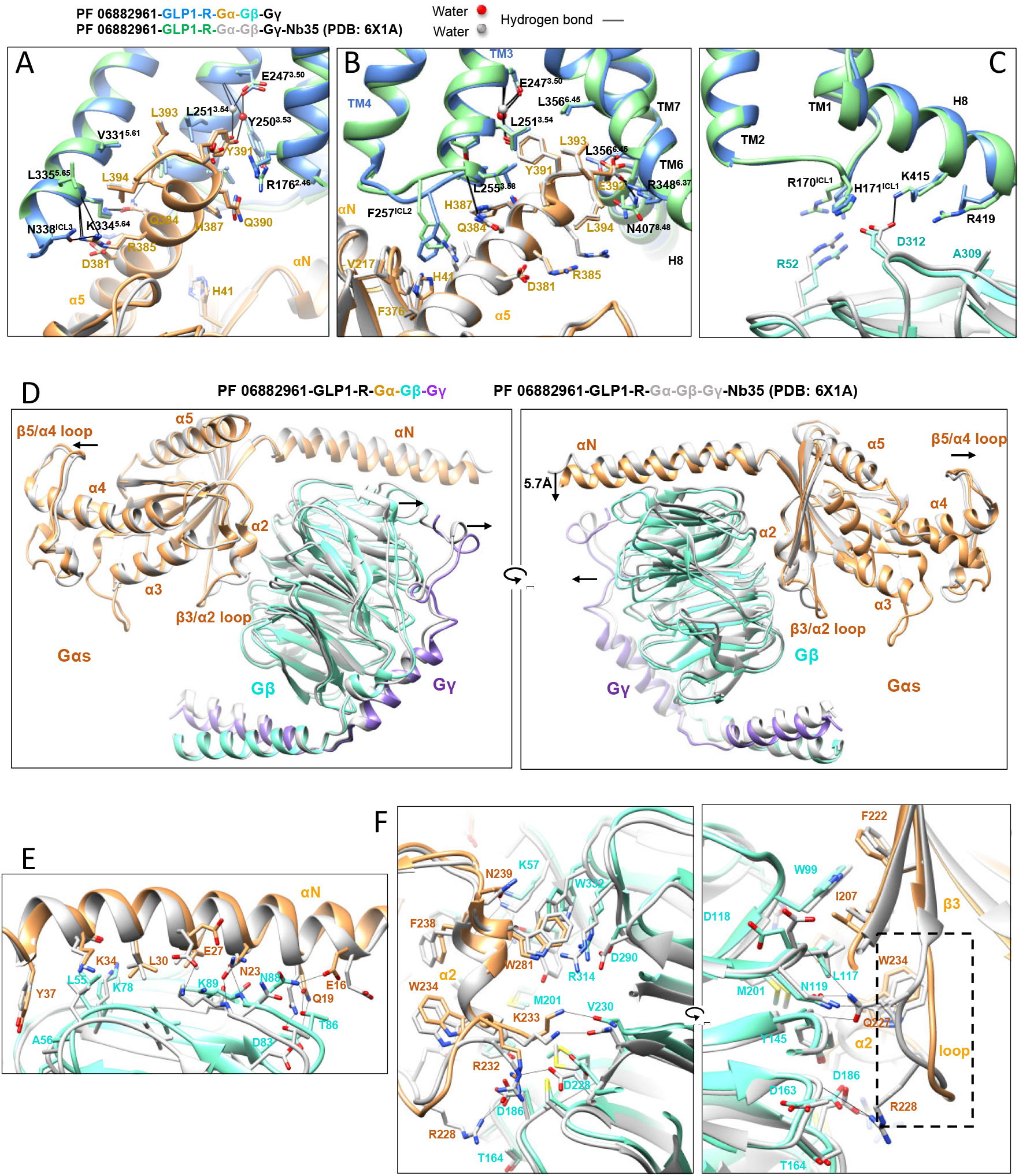
Subunit and GLP-1R interactions of the G protein heterotrimer. (**A-C**) Side views of the GLP-1R and Gs interface following superimposition of the receptor core within PF 06882961-bound complexes in the presence (PDB: 6X1A) or absence of Nb35. (**A**) Interface of Gαs and GLP-1R viewed from the intracellular portion of TM6/TM7 where TM6/TM7-H8 have been removed for clarity; (**B**) view from the intracellular portion of TM5 where TM1/2/5 have been removed for clarity; (**C**) Interface of Gβ and GLP-1R. Interacting amino acids are displayed in stick format coloured by heteroatom with protein segment colouring as noted on the figure panel. (**D**) Comparison of the orientation of heterotrimeric Gs in GLP-1R complexes formed the presence or absence of Nb35, aligned on the receptor core. Colouring denotes protein components as highlighted on the figure panels. Arrows indicate the direction of notable changes relative to 6X1A. (**E-F**) Interfaces between Gαs and Gβ subunits. Superimposition on the Gαs core, reveals differences in the interface of the αN helix and Gβ (**E**) and the interface of β_2_/β3/α2 cluster and Gβ (**F**) within PF 06882961-bound complexes in the presence (PDB: 6X1A) or absence of Nb35. The difference in the β3/α2 loop is highlighted using a dashed rectangle.

### Interactions between heterotrimeric Gs subunits

The ras-like domain of Gαs, including the αN helix, β_2_/β3/α2 cluster and the loop between α3 and β5, formed extensive interactions with the Gβ1 subunit (**Fig. 3D; Table S3**). The N-terminal α-helix of the Gαs ras-like domain primarily formed hydrogen bonds with Gβ1 (E16^αN^ with N88^β^, Q19^αN^ with D83^β^/T86^β^/N88^β^, N23^αN^ with the backbone of K89^β^), while the rest of the αN helix formed van der Waals interactions with Gβ1 (**Fig. 3E; Table S3**). In addition, the β_2_/β3/α2 cluster of the ras-like domain formed extensive hydrophobic contacts with the base of the β-propeller of the Gβ1 subunit (**Fig. 3F; Table S3**). Generally, where the residue side chain rotamers were well supported by cryo-EM density (**Suppl. Fig. 2**), there was a similar interaction pattern for most of the Gα/Gβ interface relative to the structure stabilized by Nb35, albeit with variance in some of the side chain orientations and interaction properties, including those of E16^αN^, E27^αN^ and R232^β3/α2_loop^ of Gαs (**Fig. 3E, 3F; Table S3**). In the absence of Nb35, the electron density of the β3/α2 loop of Gαs was poor even following the G protein-focussed refinement; this region was only modelled at the backbone level and the specific interactions within this region could not be defined. In contrast, the β3/α2 loop was well resolved in the Nb35-stabilized structure, in which Nb35 bridged the Gα/β interface and formed close contacts with the β3/α2 loop via both hydrophobic and polar interactions, leading to a more stable β3/α2 loop (**Suppl. Fig. 6; Table S4**). Almost the whole of the Gγ2 chain closely engages with the N-terminal helices and blades 1/4/5/6/7 of the Gβ1 subunit, via both polar contacts and hydrophobic interactions (**Fig. 3D; Table S5**), adopting a similar position to that in the Nb35-stabilised structure but with fewer interactions compared to either the Nb35-stabilised PF 06882961- or GLP-1-bound structures (**Table S5**).

Superimposition on the GLP-1R in the Nb35-stabilised and absent structures revealed a similar architecture of the heterotrimeric Gs protein, but there were differences in the orientations of each subunit relative to the receptor core (**Fig. 3D**). Compared to the Nb35-stabilized PF 06882961 structure, the αN helix of the ras-like domain in the absence of Nb35 moved towards Gβγ by approximately 6 Å (**Fig. 3D**). Interestingly, the location of the αN helix in the consensus structures also varied among different Nb35-stabilized GLP-1R-Gs complexes (**Suppl. Fig. 7**) as well as between different class B GPCR structures^15^. In addition, the β5/α4 loop that is located at the far end of Gαs was displaced away from the Gs core, relative to the Nb35-bound structure, whereas little difference was observed for the position of the α5 helix. Concurrently, in the absence of Nb35, both Gβ1 and Gγ2 were shifted away from the Gα/β interface, leading to a more stretched conformation of the heterotrimeric G protein (**Fig. 3D**).

### Dynamic analysis of PF 06882961-GLP-1R-DNGs in the absence of Nb35

An equivalent 3D variance analysis to that used to study GLP-1R complexes stabilised by Nb35^14^ was performed to understand and visualize the dynamics of PF 06882961-GLP-1R-DNGs complex without Nb35 bound. The top three principal components (modes) were extracted from the refined particle stack, with each mode displaying a specific type of variability, and the three major components of both PF 06882961-bound complexes in the presence or absence of Nb35 were recorded side by side in **Video S1**. Despite similar twisting and rocking motions of the GLP-1R relative to Gs in both complexes, notable distinctions were observed in dynamics of the heterotrimeric G protein (**Video S1, components 1 and 2**). In the absence of Nb35, a rotational motion of Gs relative to the GLP-1R was observed. In particular, the αN helix of Gαs and Gβγ dimer underwent a larger motion than other regions of Gαs, with the N-terminal portion of Gα and Gβ, as well as almost the entire Gγ subunit displaying attenuated EM density in the dynamic states, indicative of high mobility in these regions. In contrast, the G protein heterotrimer (with the exception of AHD) in the presence of Nb35 was relatively stable, despite slight motions on the protein surface (**Video S1, components 1 and 2**). At lower contour level within component 3, density for the AHD could be observed, and this domain maintained a diffuse predominantly closed, position. This contrasted with the 3D variance data from the Nb35-bound complex where the AHD underwent translational motion between open and more closed conformations relative to the Gαs ras-like domain (**Video S2**)^14^. Further 3D classification in RELION was subsequently performed, using a generous mask to include the AHD, to enable further analysis of the particle distributions in different classes. 181K particles (~32% of the whole particle stack) exhibited density for the entire AHD in a closed conformation, and 278K (~50%) particles had partial density of the AHD also in a closed conformation, while 104K (~18%) particles lacked density for the AHD (**Suppl. Fig. 8A**). The best resolved class of 181K particles was selected for an additional G protein-focussed refinement (**Fig. 1A**). Although the AHD was still poorly resolved with local resolution lower than 4 Å (**Fig. 1A**), the backbone of the AHD in the “closed” orientation that was modelled in the PF 06882961-GLP-1R-Gs-Nb35 structure (PDB: 6X1A) could be rigid body fitted well into the local refined map, demonstrating conservation of the closed conformation with that observed in the Nb35-stabilized structure (**Suppl. Fig. 8B**).

### Comparison of PF 06882961-GLP-1R-DNGs complexes imaged with 200kV and 300kV cryo-EM

The latest K3 and Falcon 4 direct electron detectors have provided improvements in the rate of image acquisition and quality of collected data, contributing to the enhanced resolutions being achieved for cryo-EM, including GPCR complexes. The potential impact of these technologies on imaging with 200kV instruments has not yet been explored. As such, we performed additional data acquisition and analysis on the PF 06882961-GLP-1R-DNGs complex as an exemplar of relevance to the potential use of 200kV imaging to GPCR drug development.

Two additional data sets were collected using either a Glacios 200kV cryo-EM or Krios 300kV cryo-EM equipped with a Falcon 4 detector and these were compared to the data described above that was collected on a Krios-K3 system. Vitrified samples imaged on either the Krios (**Suppl. Fig. 9A**) or Glacios (**Suppl. Fig. 9E**), revealed well-defined secondary structure following 2D classification (**Suppl. Fig. 9B, 9F**), with consensus 3D reconstruction of 2.8 Å and 3.2 Å, respectively, for the Krios (**Suppl. Fig. 9C**) and Glacios (**Suppl. Fig. 9G**) data sets at gold standard FSC 0.143. Local resolution in the receptor was improved by local refinement with a mask on the receptor. Similar to the original data set, there was substantial preferred orientation but good low frequency angular coverage (**Suppl. Fig. 9D, 9H**). The local resolution for the Krios-Falcon 4 data set was similar to that for the Krios-K3 data (**Fig. 4A c.f. Fig. 1A**) and 0.2 to 0.4 Å lower for the Glacios data set (**Fig. 4B**). All data sets enabled confident rotamer placement of the sidechains for most of the complex including the TM helices and binding pocket (**Suppl. Fig. 10; Fig. 4C, 4D**). While the global resolution was lower for the Glacios data, the map density in the binding site for the PF 06882961 compound was qualitatively similar to that from the maps collected with the Krios instruments, including robust density for the structural waters (**Fig. 4C, 4D; Suppl. Fig. 3A**). An overlay of PF 06882961 from the 3 structures revealed identical pose and conservation of direct interactions with the receptor, along with the water network that supports compound binding and the active conformation of the receptor (**Fig. 4E**).

**Figure 4.**
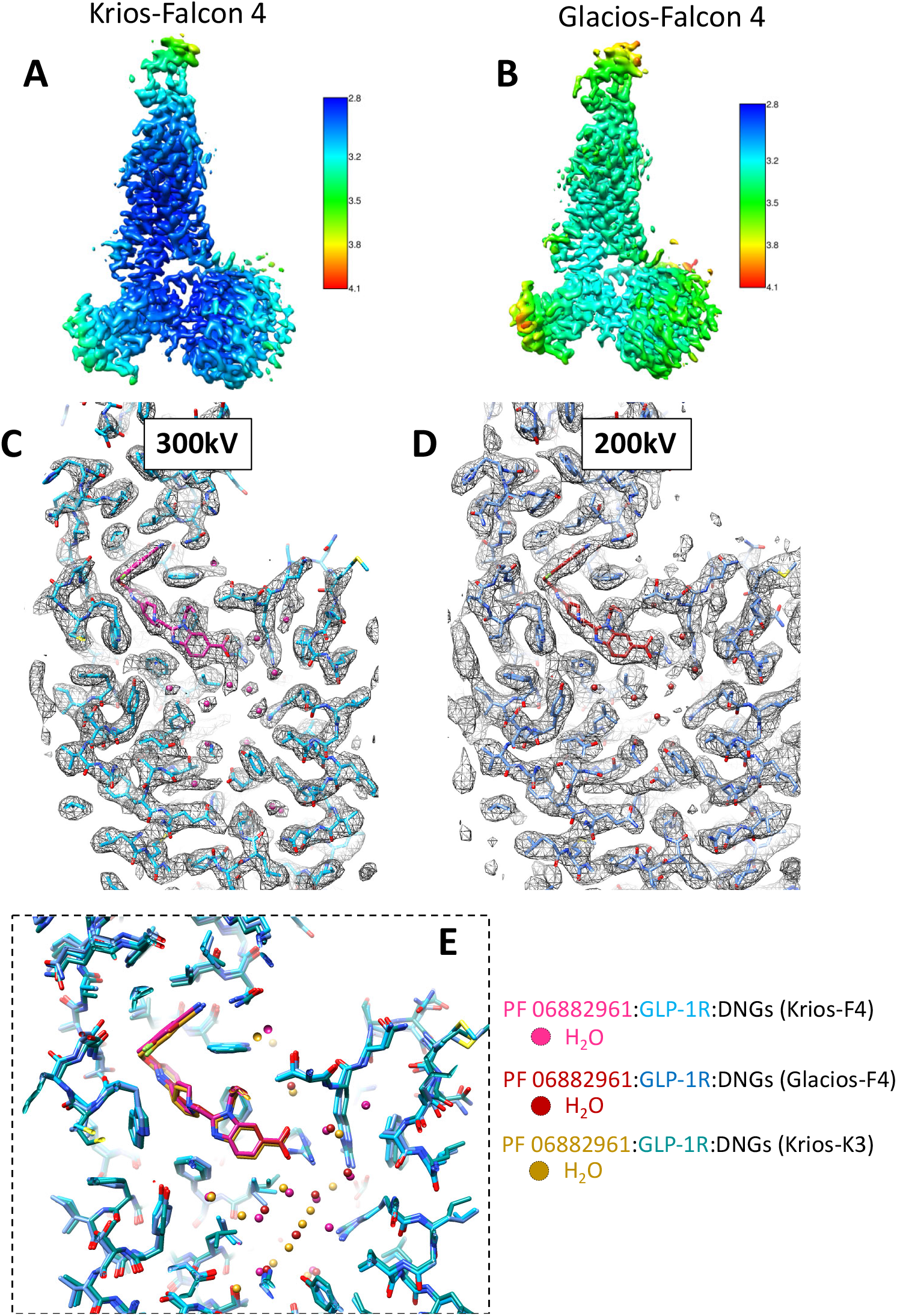
Comparison of PF 06882961-GLP-1R-DNGs complexes imaged with 200kV versus 300kV cryo-EM. (**A, B**) Local resolution-filtered EM consensus maps displaying local resolution (Å) coloured from highest resolution (dark blue) to lowest resolution (red). (**A**) Reconstruction from the Krios-Falcon 4 (F4) data set; (**B**) Reconstruction from Glacios-F4 data set. (**C**) Models of PF 06882961 (dark pink) and the GLP-1R binding cavity (sky blue) built into the receptor-focused density map for the Krios-F4 data. (**D**) Models of PF 06882961 (dark red) and the GLP-1R binding cavity (blue) built into the receptor-focused density map for the Glacios-F4 data. (**E**) Overlay of the PF 06882961-GLP-1R binding pockets for each of the structures solved in the absence of Nb35, coloured according to the displayed legend. All residues are displayed as sticks coloured by heteroatom. Waters are modelled as spheres, coloured according to the displayed legend.

## Discussion

Nb35 is a valuable tool for stabilization of GPCR-Gs complexes, leading to the crystallisation of the first Gs protein-bound GPCR active complex^6^, and has become a routine component of strategies for solution of GPCR-Gs structures by cryo-EM, either as the sole agent to enable stabilisation or in combination with other technologies including mini-Gαs^10–12^ or DNGαs^7^. The Gs protein is the canonical transducer for a large number of GPCRs^16^, and consequently structures of active-state GPCRs coupled to Gs are critical to the understanding of agonist binding to these receptors and for future drug discovery and development. Unlike Nb35 or mini-Gαs, the DNGαs technology is not protected and can be used for commercial application without restriction. In the current study, we assessed the utility of DNGαs as a stand-alone technology for stabilisation of GPCR-Gs complexes and compared this to complexes that were further stabilised by Nb35. For this analysis, we used an exemplar of current interest to the pharmaceutical industry, GLP-1R-Gs in ternary complex with the small molecule agonist, PF 06882961. Quantitatively similar yields, with similar purity and stability were obtained relative to our previously obtained complex with Nb35^14^. The global resolution of complexes solved in the absence of Nb35 was lower (2.8-2.9 Å) than that solved with Nb35 (2.5 Å), however, this was due to increased mobility of the G protein when Nb35 was absent. Of note, the local resolution for the receptor, and in particular, the PF compound binding site was equivalent for both structures, with identical poses for the ligands and interaction patterns with the receptor (**Table S1**). Importantly, the resolution in the binding pocket supported the modelling of the structural waters that play a critical role in the binding and pharmacology of PF 06882961^14^. As such, DNGαs can be used to support commercial discovery projects for agonist-GPCR-Gs complexes. However, it should be noted that this is not the only non-patented approach that could be applied for Gs complexes. The recently developed short chain antibody, scFv16, that binds across the αN of Gi/o proteins and Gβ^17,18^ has been shown to have utility for stabilisation of GPCR-G protein complexes formed with chimeric G proteins, where the Gi-αN is substituted into alternate G proteins, including stabilisation of complexes with a Gs-GiαN chimera^18^. Additional work will be required to examine the thresholds for when these different technologies alone may be sufficient for lower efficacy ligands, or when combinations of approaches may be required.

The ability to form stable complexes in the presence and absence of Nb35 also allowed us to gain insight into the impact of Nb35 on GPCR-Gs complex structure and dynamics. Broader analysis of the conformation of the receptor and of the interface between GLP-1R and the G protein heterotrimer revealed small differences in the intracellular conformation of the receptor between the two complexes, with the largest variances occurring in the conformation of ICL2 and the location of the base of TM6. It is likely that these small differences are related to the dynamic nature of GPCRs, as well as small differences associated with the conditions of vitrification and particle selection that can occur during data analysis and 3D reconstruction, rather than alterations to the mode of receptor-G protein interaction arising through Nb35 effects on the G protein. Supporting this, there were also small differences in the consensus structures between the 3 data sets collected with the PF 06882961-GLP-1R-DNGs complexes (**Suppl. Fig. 10c**). Nonetheless, the lower resolution for the G protein in each of the consensus maps for complexes in the absence of Nb35 to that solved with Nb35^14^, was indicative of higher mobility, particularly within the G protein heterotrimer, when Nb35 was absent. This was confirmed in the 3D variance analysis of the conformational dynamics of the complex comparing principal components of motion for complexes with and without Nb35, with the greatest difference observed for the relative motion between the Gα and Gβ interface. Our application of 3D variance analysis to recently solved structures of class B GPCRs, including adenomedullin-1 (AM1) and AM2 receptors^19^, the secretin receptor^20^ and GLP-1R bound to peptide and small molecule agonists^14^ has revealed common rocking and rotational motions of the G protein relative to the receptors. This is consistent with transient formation and breakage of interactions that are likely to be important in receptor-mediated G protein recruitment and activation, and low resolution for parts of ICL3 and the extension of H8 for class B GPCRs^3,8,9,15,19–28^ are consistent with transient interactions between these domains and the G protein. ICL2 exhibited densities consistent with 2 metastable conformations that exhibited differences in the consensus maps solved in the presence and absence of Nb35, and where higher resolution was apparent for the complex without Nb35. This latter observation suggests that the restrictions to the dynamics of the G protein imparted by Nb35 may impact on how the G protein engages with ICL2, a domain implicated in enabling guanine nucleotide exchange of the G protein^29^.

Agonist-bound GPCRs promote exchange of GTP for GDP to enable activation of G proteins. A key conformational change required nucleotide exchange is outward movement of the AHD of Gαs, relative to the ras-like domain. The AHD of Gαs in the nucleotide-free state is highly mobile, thereby occupying different positions around the ras-like domain^30^, and it is poorly resolved in most consensus cryo-EM maps of solved class B GPCR-G protein complexes^3,8,9,15,19–28^. When bound with Nb35, 3D multivariate analysis of PF 06882961-GLP-1R-DNGs complex revealed translational motion between open (“up” position) and more closed (“down” position) conformations of the AHD. In contrast, equivalent analysis on the no Nb35 complex resolved only a single, downward conformation of this region at relatively low resolution (**Video S2**). AHD-focussed 3D classification revealed additional insight into conformation sampling of this domain. In the absence of Nb35, the complex exhibited density encompassing the entire AHD in the “down” orientation for ~32% of the particles. However, ~50% particles had a more varied AHD conformation with only partially resolved density that nonetheless was also located predominantly in a downward orientation, while no density was resolved in 18% of particles. In the presence of Nb35, more robust density for the AHD was observed for the PF 06882961-bound complex, where three major conformations could be resolved in an “up”, “middle” or “down” orientation, with the downward conformation best resolved^14^.

Although the AHD density was not well resolved in the Nb35-absent complex, the backbone of the AHD from the PF 06882961-GLP-1R-Gs-Nb35 structure (PDB: 6X1A) could be fitted into the local refined map as a rigid body, where it occupied an identical downward position (**Suppl. Fig. 8B**). In order to visualize the AHD transition between the inactive GDP-bound conformation and the downward conformation when GLP-1R coupled, a morph between the two structures was recorded in **Video S3**. In the GDP-bound Gαs structure (PDB: 6EG8), the AHD is in close contact with the ras-like domain, forming the nucleotide binding site^31^, and it undergoes an anti-clockwise rotation to dissociate from ras-like domain upon GLP-1R coupling. A large conformational change was also observed for the α5 helix that shifts towards the receptor and rotates as the C-terminus engages within the receptor TM groove (Video S3) that is characteristic of G protein coupling; observed in Gs, Gi/o^32^ and Gq/11^33,34^ complexes with GPCRs. These changes expose the GDP binding site, facilitating dissociation of GDP that is required for GTP exchange. The observed differences in dynamics of the AHD in the absence and presence of Nb35 indicate that the relative conformational dynamics within the Gα-Gβγ interface, and indirectly the interface between the receptor and G protein, influence the associated dynamics of the AHD. While not directly comparable, structures of rhodopsin-Gt complexes have also been solved with and without an engineered form of Nb35 that can bind Gαt, where subtle differences in the orientation of Gt relative to rhodopsin were observed^35^. In that study, there were also effects on the location of the AHD observed following 3D classification consistent with an influence of the Nb on behaviour of the Gt protein albeit that both the G protein and receptor system are completely distinct from that in our study. While this new data provides additional insight into the interaction and dynamics of the Gs protein in the ternary agonist-GPCR-G protein complex there are multiple factors outside of the effect of Nb35 that may artifactually affect the dynamics of the G protein. These include distinctions in behaviour that may occur in the protein in the detergent micelle when compared with native membranes, and the impact of the mutations in the DNG protein. As such, the extrapolation of the structural and dynamic data needs to be done with caution.

In addition to the ability to form stable GPCR-Gs complexes without patented technologies, we wanted to explore the extent to which the evolution in our understanding of optimal processes for vitrification^36^, detector technology and 3D reconstruction software might enable the use of 200kV cryo-EM to support structure assisted drug discovery and development. We used a bottom mounted Falcon 4 detector with 200kV microscopy to collect data for the PF 06882961-GLP-1R-DNGs sample, and an equivalent data set was collected with this detector and a 300kV Krios as a direct comparator for the detector, and to assess imaging with the K3 versus Falcon 4 detector. In this latter comparison, similar, high-resolution data was generated indicating that both detectors perform well for 300kV imaging. A more nuanced comparison was not possible as data were collected from grids vitrified in different facilities. Excitingly, while the global resolution was 0.2-0.4 Å lower, the structure determined from the Glacios-Falcon 4 data was effectively identical for the small molecule binding pocket, including the water network within this pocket. While direct comparator data with 300kV imaging is not available, a recent publication in BioRxiv reports a CCR5-agonist-G protein complex resolved to 3.1 Å resolution using a Glacios-K3 system^37^, further supporting the potential utility of 200kV cryo-EM for GPCR structure determination.

In conclusion, our data illustrate that recent, license-free, technologies for stabilisation of agonist-GPCR-Gs complexes can yield structures with equivalent resolutions in the GPCR, including within the drug binding pocket, and which can reveal key structural waters involved in drug-receptor interactions. Moreover, resolutions that can robustly support structure-assisted GPCR drug discovery can be achieved with a 200kV cryo-EM, opening up broader application of cryo-EM within the pharmaceutical and biotechnology industries.

## Acknowledgments

The authors thank Yi-Lynn Liang for assistance with protocols for biochemistry.

## Author contributions

X.X. expressed and purified receptor complexes. R.D., I.D., L.Y. and A.K. performed cryo-sample preparation and imaging to acquire EM data. X.X., R.J., M.J.B. processed the EM data and performed EM map calculations, including 3D multivariate analysis, built the protein models for consensus maps and performed refinement. X.X., R.J., M.J.B., D.W. assisted with data interpretation, figure and manuscript preparation. All authors reviewed and edited the manuscript; M.J.B., P.M.S., and D.W. designed the project, and interpreted data. M.J.B., P.M.S., and D.W., supervised the project. X.X., M.J.B., D.W. and P.M.S. generated figures and/or wrote the manuscript.

## Data and materials availability

All relevant data are available from the authors and/or included in the manuscript or Supplementary Information. Atomic coordinates and the cryo-EM density maps have been deposited in the Protein Data Bank (PDB) under accession numbers 7LCI (Krios-K3 data set), 7LCJ (Krios-F4 data set) and 7LCK (Glacios-F4 data set), and Electron Microscopy Data Bank (EMDB) accession numbers EMD-23274 (Krios-K3), EMD-23275 (Krios-F4) and EMD-23276 (Glacios-F4).

## On-line Methods

### Constructs

For complex formation, the previously described human GLP-1R, modified to replace the native signal peptide with that of haemagglutinin (HA) and containing an N-terminal Flag tag and C-terminal 8xHis tag flanked by 3C protease cleavage sites (LEVLFQGP) was used^25^. Similarly, the previously described and characterised DNGαs was used^15,19,20^. GLP-1R, DNGαs, Gβ1 and Gγ2 were expressed in insect cells following established protocols^24^. 8xHis tagged Nb35 was provided by B. Kobilka^6^.

### Insect cell expression

The Bac-to-Bac Baculovirus Expression System (Invitrogen) was used to generate high-titre recombinant baculovirus (> 10^9^ viral particles per ml). Recombinant baculovirus was produced by transfecting recombinant bacmids (2-3 μg) into *Sf9* cells (5 mL, density of 4×10^5^ cells per mL) using FuGENE HD Transfection Reagent (Promega) and Opti-MEM Reduced Serum Media (Thermo Fisher Scientific). After 5 d incubation at 27°C, P0 viral stock was harvested as the supernatant of the cell suspension to produce high-titre viral stock (P1 and P2 virus). Viral titres were analysed by flow cytometry on cells stained with gp64-PE antibody (Expression Systems). Human GLP1R, DNGαs and Gβ1-γ2 were co-expressed by infecting *Tni* insect cells at a density of 3.5 million/mL with P2 baculovirus at multiplicity of infection (MOI) ratio of 3:3:1. Culture was harvested by centrifugation 48 h post infection and cell pellets were stored at −80°C.

### Complex formation and purification

GLP-1R-DNGs complex formation and purification were performed as described by Liang, *et al*.^14,24^. Cell pellets (from 1 L insect cell culture) were thawed and suspended in 20 mM HEPES pH 7.4, 50 mM NaCl, 5 mM CaCl_2_, 2 mM MgCl_2_ supplemented with cOmplete Protease Inhibitor Cocktail tablets (Sigma Aldrich) and benzonase (Merck Millipore). Complex was formed by adding 50 μM PF 06882961 and apyrase (25 mU/ml, NEB). The suspension was incubated for 1 h at room temperature. The complex was solubilized from membrane using 0.5% (w/v) LMNG and 0.03% (w/v) CHS (Anatrace) for 1 h at 4°C. Insoluble material was removed by centrifugation at 30,000 g for 20 min and the solubilized complex was immobilized by batch binding to M1 anti-Flag affinity resin in the presence of 5 mM CaCl_2_. The resin was packed into a glass column and washed with 20 column volumes of 20 mM HEPES pH 7.4, 100 mM NaCl, 2 mM MgCl_2_, 5 mM CaCl_2_, 5 μM agonist, 0.01% (w/v) LMNG and 0.0006% (w/v) CHS followed by elution with buffer containing 5 mM EGTA and 0.1 mg/ml FLAG peptide. The complex was then concentrated using an Amicon Ultra Centrifugal Filter (MWCO, 100 kDa) and subjected to size-exclusion chromatography (SEC) on a Superdex 200 Increase 10/300 column (GE Healthcare) that was pre-equilibrated with 20 mM HEPES pH 7.4, 100 mM NaCl, 2 mM MgCl_2_, 5 μM agonist, 0.01% (w/v) MNG and 0.0006% (w/v) CHS to separate complex from contaminants. Eluted fractions consisting of receptor and G protein complex were pooled and concentrated to 5 mg/mL. The complex samples were flash frozen in liquid nitrogen and stored at −80°C.

### SDS-PAGE and Western blot analysis

Samples from important steps during purification were collected and analysed by SDS–PAGE and western blot. TGX™ Precast Gel (BioRad) was used to separate proteins within samples at 200 V for 30 min. Subsequently, gels were either stained by Instant Blue (Sigma Aldrich) or immediately transferred to PVDF membrane (BioRad) at 100 V for 1 h. The proteins on the PVDF membrane were probed with two primary antibodies simultaneously, rabbit anti-Gs C-18 antibody (cat. no. sc-383, Santa Cruz) against Gαs subunit and mouse poly-His antibody (cat. no. 34660, QIAGEN) against His tags. The membrane was washed and incubated with secondary IRDye anti-mouse and IRDye anti-rabbit antibodies (LI-COR). The membranes were imaged at wavelengths of 680 and 800 nm to visualize His-tagged proteins (GLP-1R, Gβ1 and Nb35) and Gαs, respectively, on a Typhoon 5 imaging system (Amersham).

### Negative staining and data processing

The GLP-1R complex samples were diluted to 0.006 mg/mL in 20 mM HEPES pH 7.4, 100 mM NaCl, 2 mM MgCl_2_ and 5 μM agonist and applied to continuous carbon grids (EMS). The grids were stained with 0.8% (w/v) uranyl formate solution and imaged using a cryo-EM Tecnai™ T12 TEM at 120 kV. Around 50 images were collected with a magnified pixel size of 2.06 Å, and ~20,000 particles were auto picked, extracted and 2D classified by RELION-3.1-beta.

### Vitrified sample preparation and data collection

3 μL of the sample was applied to glow-discharged UltrAuFoil R1.2/1.3 Au 300 gold foil grids (Quantifoil GmbH, Großlöbichau, Germany) and were flash frozen in liquid ethane using a Vitrobot mark IV (Thermo Fisher Scientific, Waltham, Massachusetts, USA) set at 100% humidity and 4°C for the preparation chamber and 10 s blot time. Cryo-EM data was collected on a Titan Krios G3i microscope (Thermo Fisher Scientific) operated at an accelerating voltage of 300 kV with a 50 μm C2 aperture at an indicated magnification of 105 kX in nanoprobe EFTEM mode. A Gatan K3 direct electron detector positioned post a Gatan BioQuantum energy filter (Gatan, Pleasanton, California, USA), operated in a zero-energy-loss mode with a slit width of 15 eV was used to acquire dose fractionated images of the samples with a 100 μm objective aperture. Movies were recorded in hardware-binned mode (previously called counted mode on the K2 camera) with the experimental parameters listed in **Table S6** using a 9-position beam-image shift acquisition pattern by custom scripts in SerialEM^38^.

### Cryo-EM data processing for Krios-K3 data

The cryo-EM data workflow is illustrated in **Suppl. Fig 11**. 5508 movies of PF 06882961-GLP-1R-DNGs complex were motion corrected using MotionCor2^39^ and subjected to CTF estimation with GCTF software^40^. 4.3 million particles were picked from corrected micrographs using crYOLO software^41^. Picked particles were extracted and 2D classified using RELION (version 3.1). 1.8 million particles were selected and used to generate the initial 3D model by using the SGD algorithm and subsequently subjected to 3D classification. The best-looking class of 1.1 million particles was subjected to Bayesian particle polishing and another round of 2D classification. The resulting set of 563K particles was subjected to three rounds of CTF refinement and 3D auto refinement, followed by another round of 3D classification with fine angular sampling and 3D refinement that produced a 3D map at 3.4 Å in a 288 pixels box (0.83 Å/pixel) from a set of 386K particles. The particle set was then subjected to 3D refinement and post processed with a mask excluding the detergent micelle and Gαs α-helical domain (AHD), resulting in the final consensus map at 2.9 Å (**Suppl. Fig. 11A**). Local resolution was determined using RELION with half-reconstructions as input maps.

The receptor was further refined by using a loose mask of PF 06882961-bound receptor alone in RELION. To improve the electron density of G protein, particularly the interface to the solvent, the refined particle stack was subjected to 3D classification without alignment using a general mask of the G protein excluding AHD, from which the best-looking class of 134K particles with clear electron density of each subunit of the heterotrimeric Gs protein was selected, followed by local refinement focused on G protein using a tighter mask of Gs excluding AHD (**Suppl. Fig. 11B**). To improve the resolution of AHD, the refined particle stack (563K particles) was subjected to 3D classification without alignment into 4 classes using a broad mask of the AHD. Each particle class (181K, 93K, 185K or 104K) was 3D auto-refined, and the best resolved class of 181K particles was further refined and post processed with a G protein mask to generate a final map including the AHD (**Suppl. Fig. 11B**).

### Atomic model refinement

The model of PF 06882961-GLP-1R-Gs-Nb35 (PDB: 6X1A) was used as initial template for modelling. The template was fit into the cryo-EM density map with the MDFF routine in NAMD^42^. The fitted models were further refined by manual model building in COOT^43^ and real space refinement in PHENIX based on the consensus maps and receptor-focused maps^44^. The G protein model was further optimized according to the G protein-focused map. Cryo-EM density for residues between S115^ECD^ and R117^ECD^ in the ECD region was discontinuous and these residues were omitted from the final model. The final models were subjected to global refinement and comprehensive validation as implemented in PHENIX. The cryo-EM data collection, refinement and validation statistics for all complexes are reported in **Table S6**.

### Model residue interaction analysis

Interactions in the PDB files between the bound agonist and the receptor were analyzed using the “Dimplot” module within the Ligplot^+^ program (v2.2)^45^. Hydrogen bonds were additionally analysed using the UCSF Chimera package^46^, with relaxed distance and angle criteria (0.4 Å and 20 degree tolerance, respectively).

### Cryo EM dynamics analysis

3D variability analysis implemented in cryoSPARC (v2.9)^47^ was performed to understand and visualize the dynamics in GLP-1R complexes, as previously described for analysis of the dynamics of adrenomedullin receptors^19^. The particle stack from the RELION consensus refinement was imported into the cryoSPARC environment. 3D refinement was performed, using a low pass filtered RELION consensus map as an initial model and a generous mask excluding the detergent micelle created in RELION as default. 3D variability was analysed across 3 principal components that accounted for the most common motions and the 20 volume frame data in each motion was generated in cryoSPARC^47^. Output files were visualized in UCSF ChimeraX volume series and captured as movies^48^.

### Methods for cryo-EM data collected with the Falcon 4 detector

#### Glacios-Falcon 4

##### Vitrified sample preparation and data collection

Samples (3 μL) were applied to a glow-discharged UltrAufoil R0.6/1 300 mesh holey grid (Quantifoil GmbH, Großlöbichau, Germany) and were flash frozen in liquid ethane using the Vitrobot mark IV (Thermo Fisher Scientific, Waltham, Massachusetts, USA) set at 100% humidity and 4°C for the prep chamber with a blot time of 10s. Data were collected on a Glacios microscope (Thermo Fisher Scientific) operated at an accelerating voltage of 200 kV with a 50 μm C2 aperture, a 100 μm objective aperture and at an indicated magnification of 150kX in nanoprobe TEM mode. A bottom mounted Falcon 4 direct electron detector operated in Electron Event Representation (EER) mode was used to acquire images of the samples. Movies were recorded in super-resolution mode with an exposure time of 7.16 seconds amounting to a total dose of 55.8 e-/Å^2^ at a dose rate of 7 e-/pixel/second. Defocus range was set between −0.5 to −1.5 μm using 0.2 μm increments.

##### Data processing

10307 micrographs were motion corrected using the RELION 3.1 EER motion correction package with binning to a physical pixel size of 0.95 Å/pixel. Dose weighted averaged micrographs then had their CTF parameters estimated using CTFFIND 4.1.14 and were picked using the RELION Laplacian of Gaussian picker to yield 5.62 M particles. Poorly picked particles were filtered by successive rounds of 2D and 3D classification and a homogeneous particle stack was subjected to 3D refinement and particle polishing as implemented in RELION 3.1. A further round of 2D classification was performed to yield a particle stack of 1.13 M particles which underwent a further round of 3D refinement then CTF envelope refinement. A 3D classification with local Euler angle search was performed to subclassify the particles that led to the most well resolved 3D class comprising of 632 k particles. These particles went onto the final consensus 3D refinement (3.24 Å (FSC=0.143)) and a focussed 3D refinement on the receptor portion of the complex (3.28 Å (FSC-0.143)). The data processing workflow is summarised in **Suppl. Fig. 12B**. The focussed refinement maps were used for model fitting of the receptor and ligand. This was achieved by a real space refinement of the model generated from the Titan-K3 data using the PHENIX software package. Waters were manually modelled followed by manual model adjustment in COOT followed by a final real space refinement.

#### Titan Krios-Falcon 4

##### Vitrified sample preparation and data collection

Samples (3 μL) were vitrified as described for the Glacios data collection. Data were collected on a Titan Krios G4 microscope (Thermo Fisher Scientific) operated at an accelerating voltage of 300 kV with a 50 μm C2 aperture at an indicated magnification of 96kX in nanoprobe TEM mode. A bottom mounted Falcon 4 direct electron detector operated in Electron Event Representation (EER) mode was used to acquire images of the samples. Movies were recorded in super-resolution mode with an exposure time of 7 seconds amounting to a total dose of 60 e-/Å^2^ at a dose rate of 5.77 e-/pixel/second. Defocus range was set between −0.5 to −1.5 μm using 0.2 μm increments.

##### Data processing

6450 micrographs were processed in a similar manner as described for the 200 kV dataset with binning to a physical pixel size of 0.82 Å/pixel during motion correction. The major difference was that after a consensus refinement in RELION 3.1 the particle stack was imported into CRYOSPARC (version 3.0) and subjected to a further round of 2D classification. The most well resolved classes were then underwent a consensus refinement using the Non-Uniform 3D refinement as implemented which led to both an improvement in map resolution (2.82 Å (FSC=0.143)) and overall map quality. The data processing workflow is summarised in Suppl. Fig. 12A. A model of the receptor was also modelled in a similar manner as described for the 200 kV dataset.

