## Supplemental Figures and Tables for "Evolving cryo-EM structural approaches for GPCR drug discovery"

Supplementary Figures 1 – 12.
Supplementary Video Legends 1 – 3.
Supplementary Tables 1 – 6.

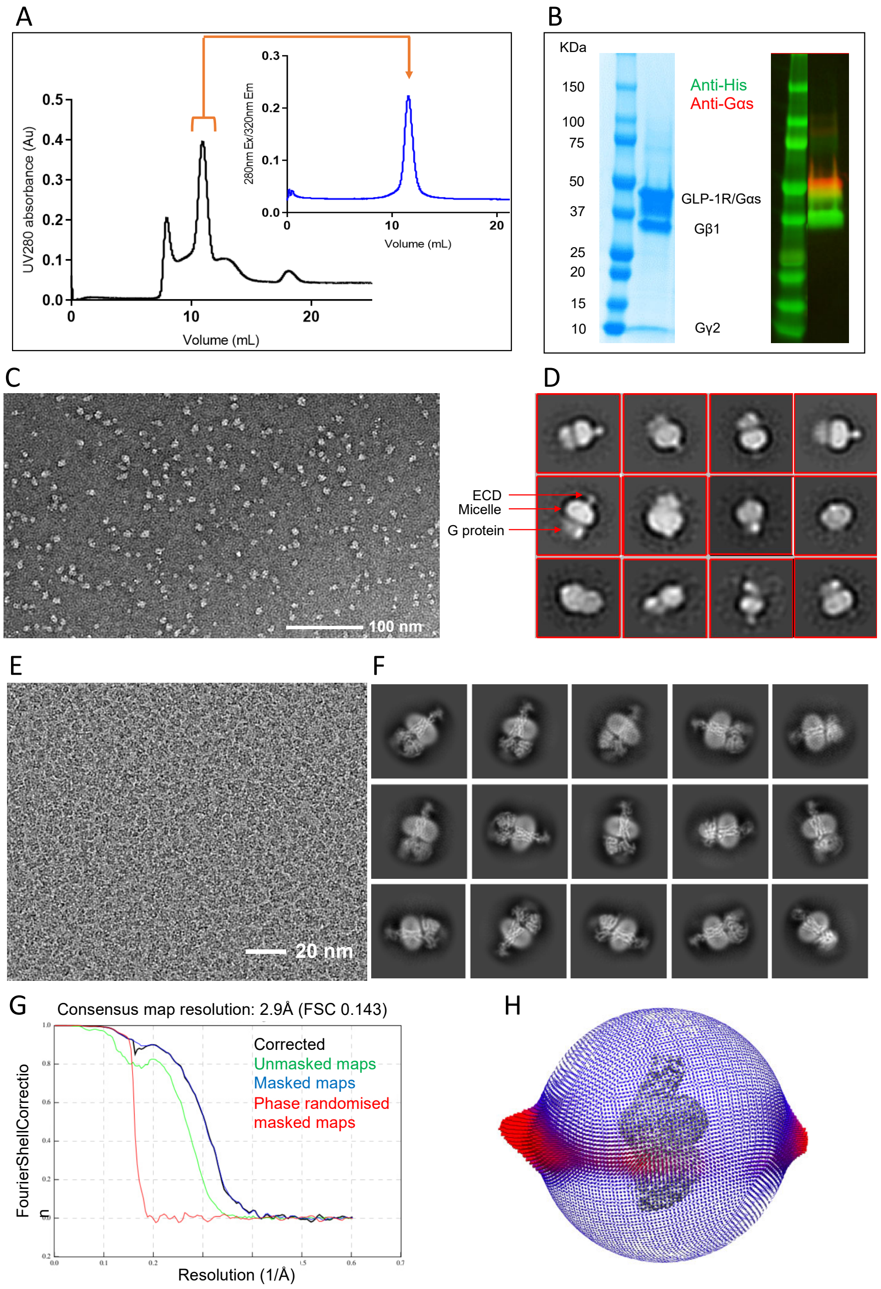


**Supplementary Figure 1. Purification and characterisation of the PF 06882961-GLP-1R-Gs complex.** (**A**) Size exclusion chromatography (SEC) profile of the complex, post-FLAG affinity purification. The peak containing the complex is highlighted by the orange bracket. The peak fractions were pooled and concentrated with purity confirmed by SEC (inset); (**B**) SDS–PAGE/Coomassie blue stain (left) and western blot of the purified complex sample illustrating that all expected components were present (right). Anti-His antibody detects GLP-1R-His, and Gβ-His (green) and anti-Gs antibody detects Gαs (red); (**C**) Representative TEM micrograph of complex particles embedded in negative stain; (**D**) Representative 2D class averages of complex particles from negative stain TEM, confirming stability of the ternary complexes. (**E**) Exemplar micrograph and (**F**) 2D class averages of cryo-EM projections of the complex; (**G**) Gold standard Fourier shell correlation 0.143 (FSC) curves for the final consensus maps and map validation from half maps, showing the overall nominal global resolution; (**H**) 3-D histogram representation of the Euler angle distribution of all the particles used in the reconstruction overlaid on the density map illustrated on the same coordinate axis.

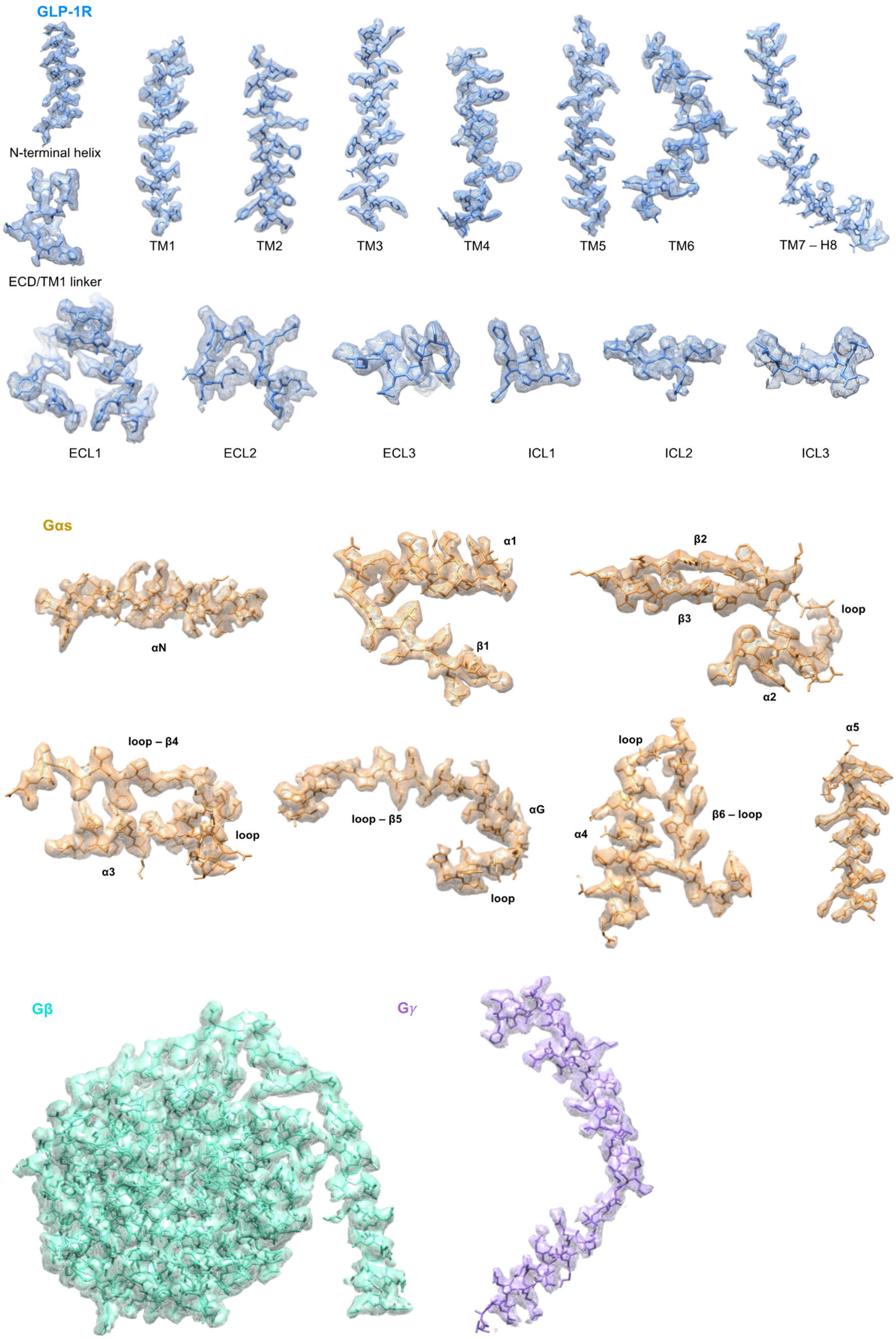


**Supplementary Figure 2. Representative atomic modelling into maps of the PF 06882961-GLP-1R-Gs complex.** Models of all seven TM helices, ECLs and ICLs of the GLP-1R (blue), selected elements of the Gαs ras-like domain (gold), Gβ (cyan) and Gγ (purple) are illustrated in the density map (mesh) generated from the receptor-focused map (GLP-1R) or G protein-focused map (Gα, Gβ, Gγ) via zone of 2 Å and mask on each component using UCSF ChimeraX. All residues are displayed in stick format.

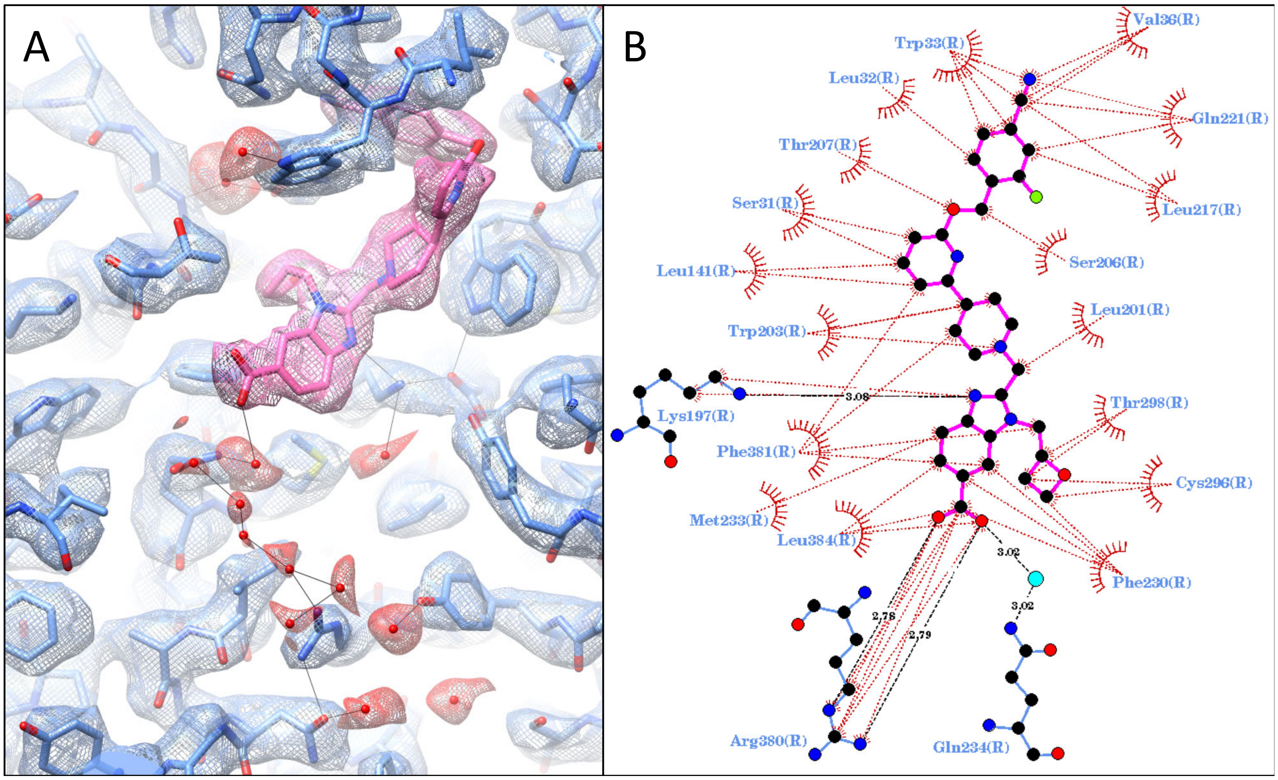


**Supplementary Figure 3. Interactions of PF 06882961 within the binding cavity of the GLP-1R.** (**A**) Models of PF 06882961 (pink) and the GLP-1R binding cavity (blue) built into the receptor-focused density map. All residues are displayed as sticks coloured by heteroatom. Waters are modelled as red spheres. (**B**) Interactions between PF 06882961 and GLP-1R as determined by Ligplot+[45]. Hydrophobic interactions are illustrated by red lines between PF 06882961 (pink stick format) and GLP-1R residues (red arcs, blue residue labels). Amino acids involved in hydrogen bonds are shown in atomic detail with hydrogen bonds shown as dashed black lines. Blue spheres indicate waters that directly interact with the ligand.

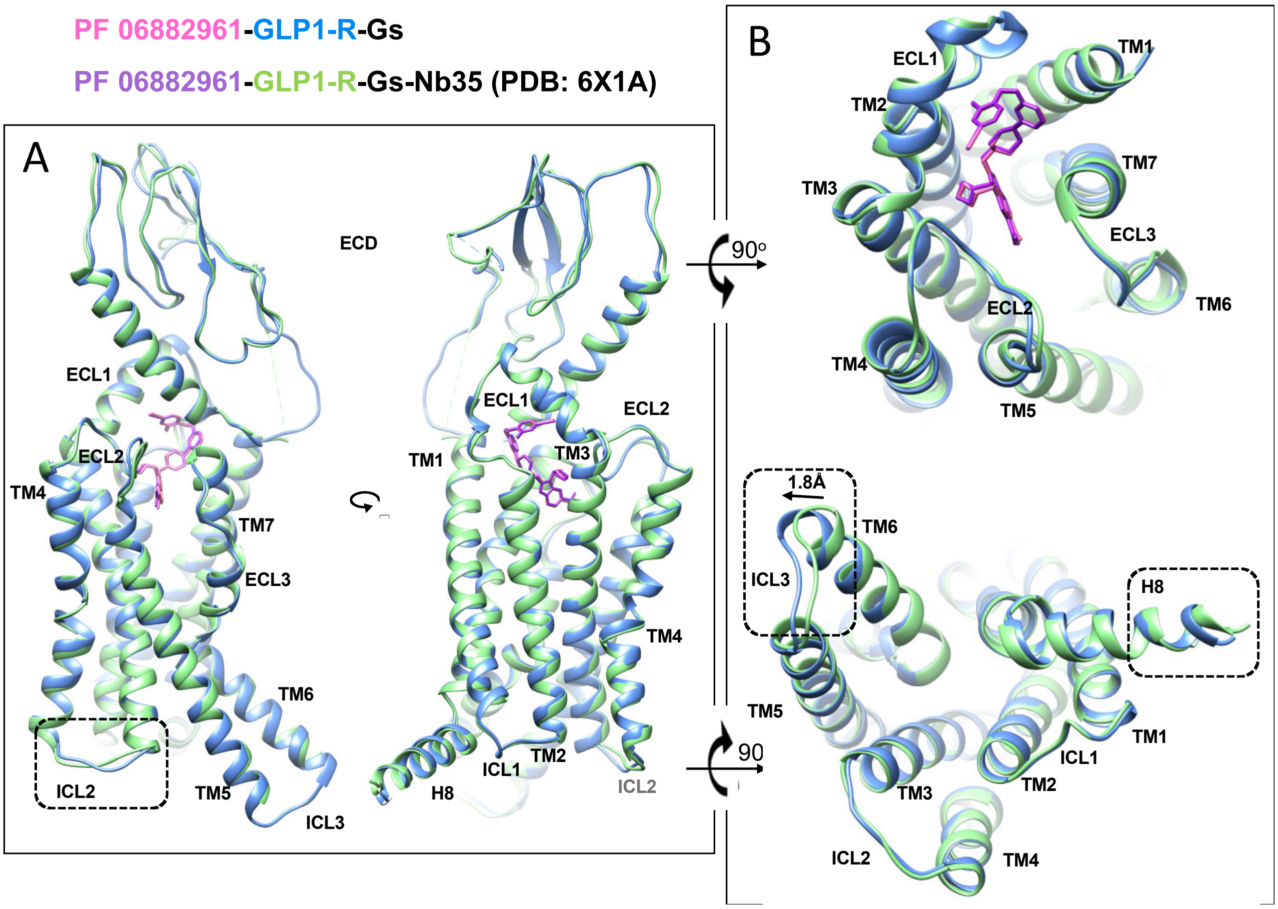


**Supplementary Figure 4. Comparison of GLP-1R conformations in the presence or absence of Nb35.** (**A**) Superimposition of the PF 06882961-bound GLP-1R in the absence of Nb35 and the Nb35-stabilized structure (PDB: 6X1A) from side view; (**B**) extracellular view (top) and intracellular view (bottom). PF 06882961 and the receptor are displayed in stick and ribbon format, respectively. Colouring denotes different components as highlighted on the figure panel. Notable differences are highlighted using dashed rectangles, with arrows showing the direction of notable changes relative to 6X1A. The distance between the intracellular tips of TM6 was 1.8 Å when measured at Cα of T343$^{6.32}$.

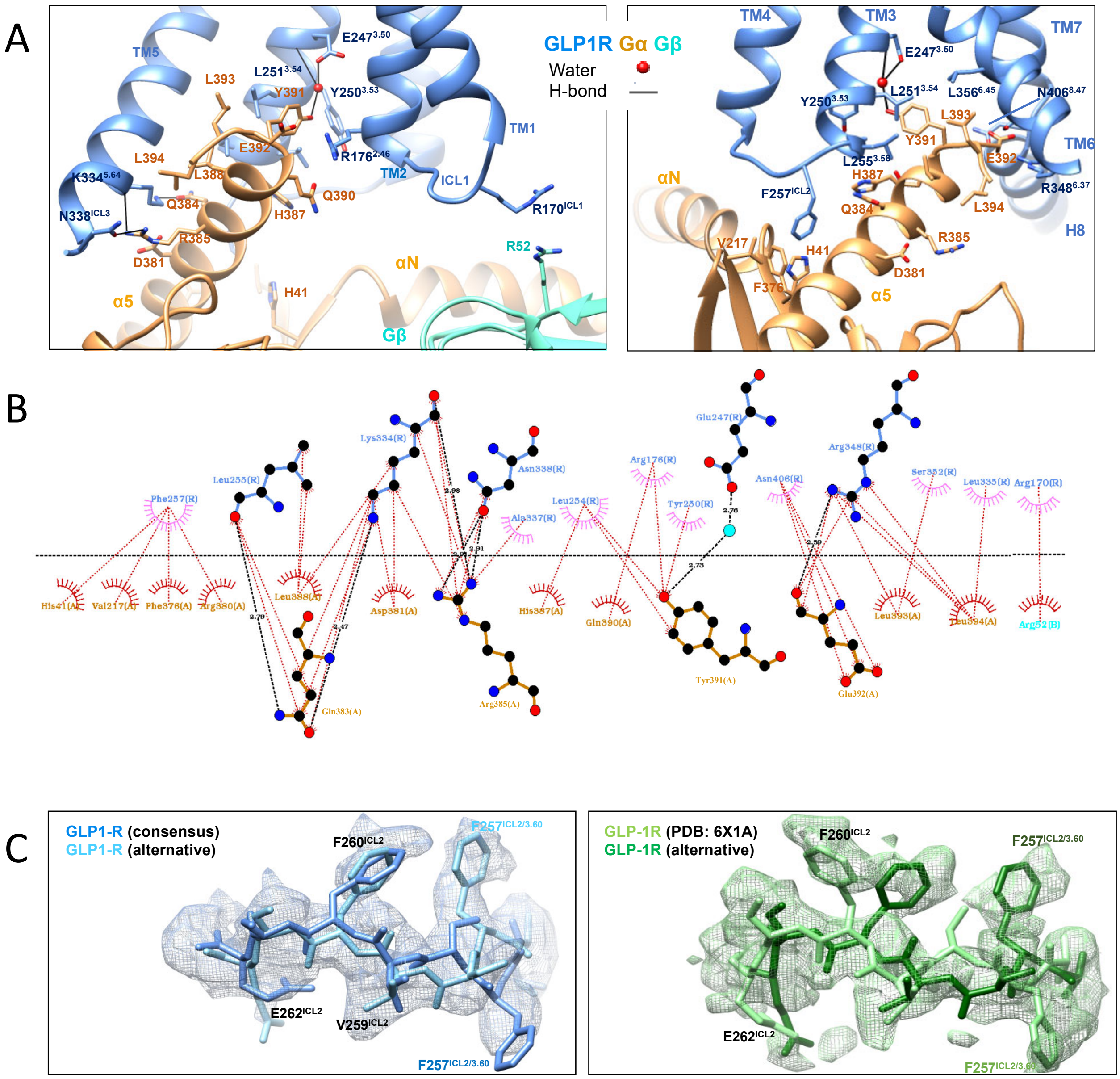


**Supplementary Figure 5. Interface between GLP-1R and G protein.** (**A**) Side view of the GLP-1R-G protein interface. Left: viewed from the intracellular portion of TM6/TM7 where TM6/TM7-H8 have been removed for clarity; Right: viewed from the intracellular portion of TM5 where TM1/2/5 have been removed for clarity. Black lines depict hydrogen bonds as determined using UCSF Chimera. Residues involved in direct or water-mediated (red sphere) interactions are displayed in stick format coloured by heteroatom, with the backbone in ribbon format. Colouring denotes different components as highlighted on the figure panels. (**B**) Interactions between GLP-1R and Gs as determined by Ligplot+. GLP-1R residues are located above the dashed black line, and G protein residues below the line. Hydrophobic interactions are illustrated by red (Gs) or pink (GLP-1R) arcs, and interacting residues are joined by a red line. Amino acids involved in hydrogen bonds are shown in atomic detail with hydrogen bonds shown as dashed black lines. Blue spheres indicate waters that directly interact with Gs; (**C**) Two ICL2 models were independently built into the electron density (mesh) in the absence (left) or presence of Nb35 (right). Residue side chains within ICL2 are highlighted in stick format with different colours for the consensus or alternate conformations (colouring theme as displayed on the figure panels).

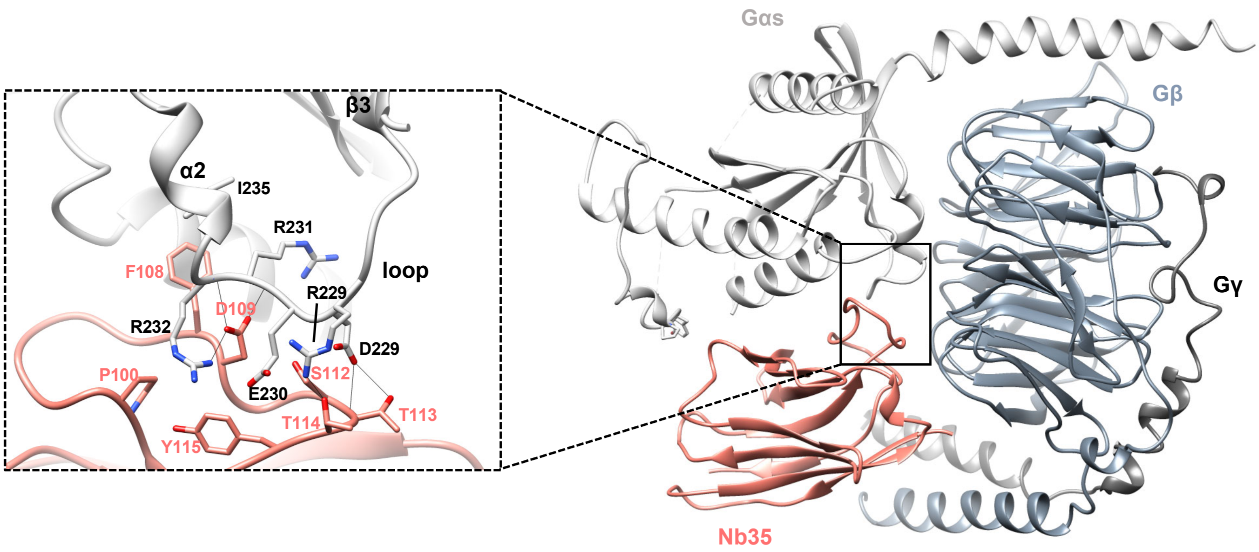


**Supplementary Figure 6. Nb35 stabilizes the interface between Gαs and Gβ.** Nb35 (salmon) binds within the interface of Gαs (light grey) and Gβ (slate grey), with no contact with Gγ (dark grey) (right). The interface of Nb35 and the β3/α2 loop of Gαs is highlighted in the dashed rectangle (left). Proteins are displayed in ribbon format. Residues involved in interactions are displayed in stick format coloured by heteroatom, with the backbone in ribbon format.

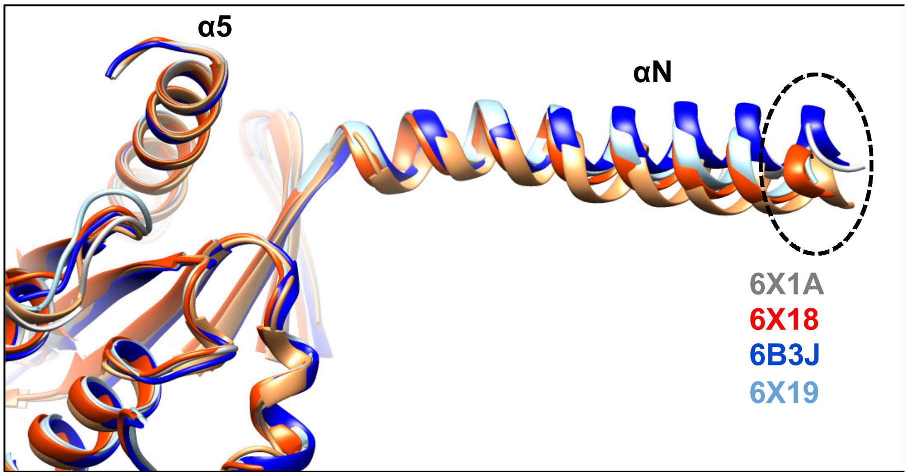


**Supplementary Figure 7. Variations in the orientation of the αN helix of Gαs for GLP-1R-Gs complex structures.** Overlay of Gαs in Nb35-stabilized GLP-1R structures bound with different ligands, including GLP-1 (PDB: 6X18, red), PF 06882961 (PDB: 6X1A, grey), CHU-128 (6X19, light blue) and Exendin P5 (PDB: 6B3J, blue), as well as the PF 06882961-bound Nb35-absent structure (orange). Notable differences in the location of the Gαs N terminus have been highlighted using a dashed oval.

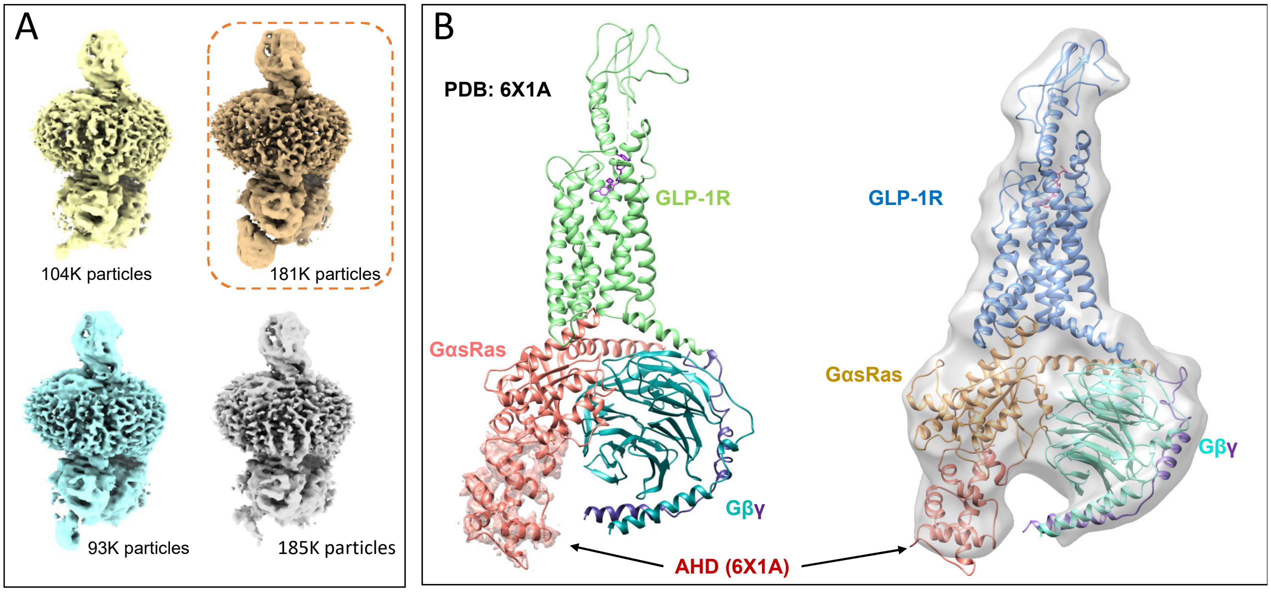


**Supplementary Figure 8. 3D classification of AHD orientations in the PF 06882961-GLP-1R-DNGs complex.** (**A**) AHD-focused 3D classification in RELION reveals different particle contribution to four classes. One class of 181K particles contains density for the entire AHD in the "down" position; two classes of 93 and 185K particles have partial densities; another class containing 104K particles lack AHD density. The best resolved class is highlighted using a dashed rectangle in orange; (**B**) The AHD density from the best resolved classes in 6X1A (+Nb35) (left panel), and the PF 06882961-GLP-1R-DNGs complex (right panel). The backbone of the AHD of 6X1A (salmon) was rigid-body fitted to the density in the "Nb35 absent" map (transparent grey surface).

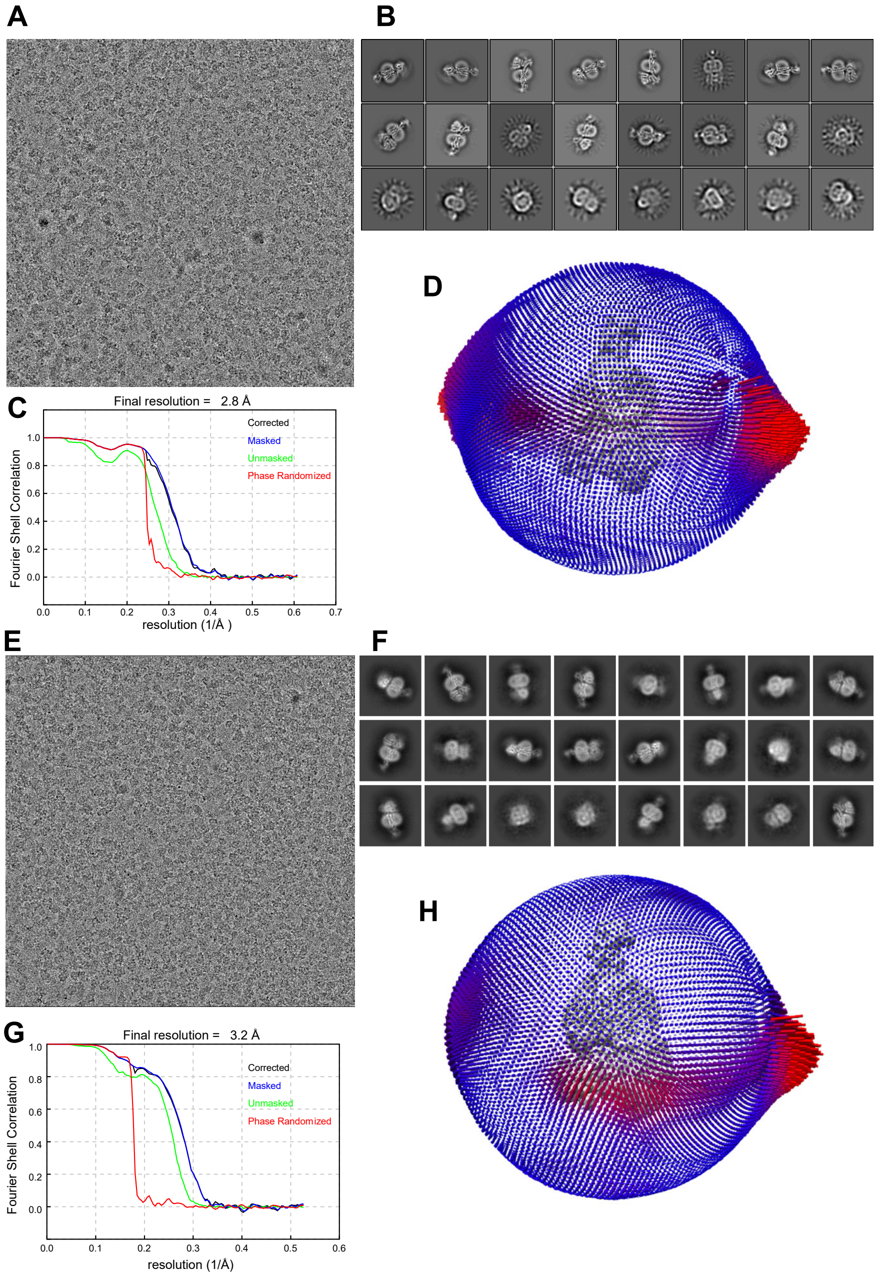


**Supplementary Figure 9. Cryo-EM structure determination of PF 06882961-GLP-1R-DNGs complexes using 300kV (Krios-Falcon 4) or 200kV (Glacios-Falcon 4) imaging.** (**A-D**) Data collected using the 300kV Krios and Falcon 4 detector. (**A**) Exemplar micrograph and (**B**) 2D class averages of cryo-EM projections of the complex; (**C**) Gold standard Fourier shell correlation 0.143 (FSC) curves for the final consensus maps and map validation from half maps, showing the overall nominal global resolution; (**D**) 3-D histogram representation of the Euler angle distribution of all the particles used in the reconstruction overlaid on the density map illustrated on the same coordinate axis. (**E-H**) Data collected using the 200kV Glacios and Falcon 4 detector. (**E**) Exemplar micrograph and (**F**) 2D class averages of cryo-EM projections of the complex; (**G**) Gold standard Fourier shell correlation 0.143 (FSC) curves for the final consensus maps and map validation from half maps, showing the overall nominal global resolution; (**H**) 3-D histogram representation of the Euler angle distribution of all the particles used in the reconstruction overlaid on the density map illustrated on the same coordinate axis.

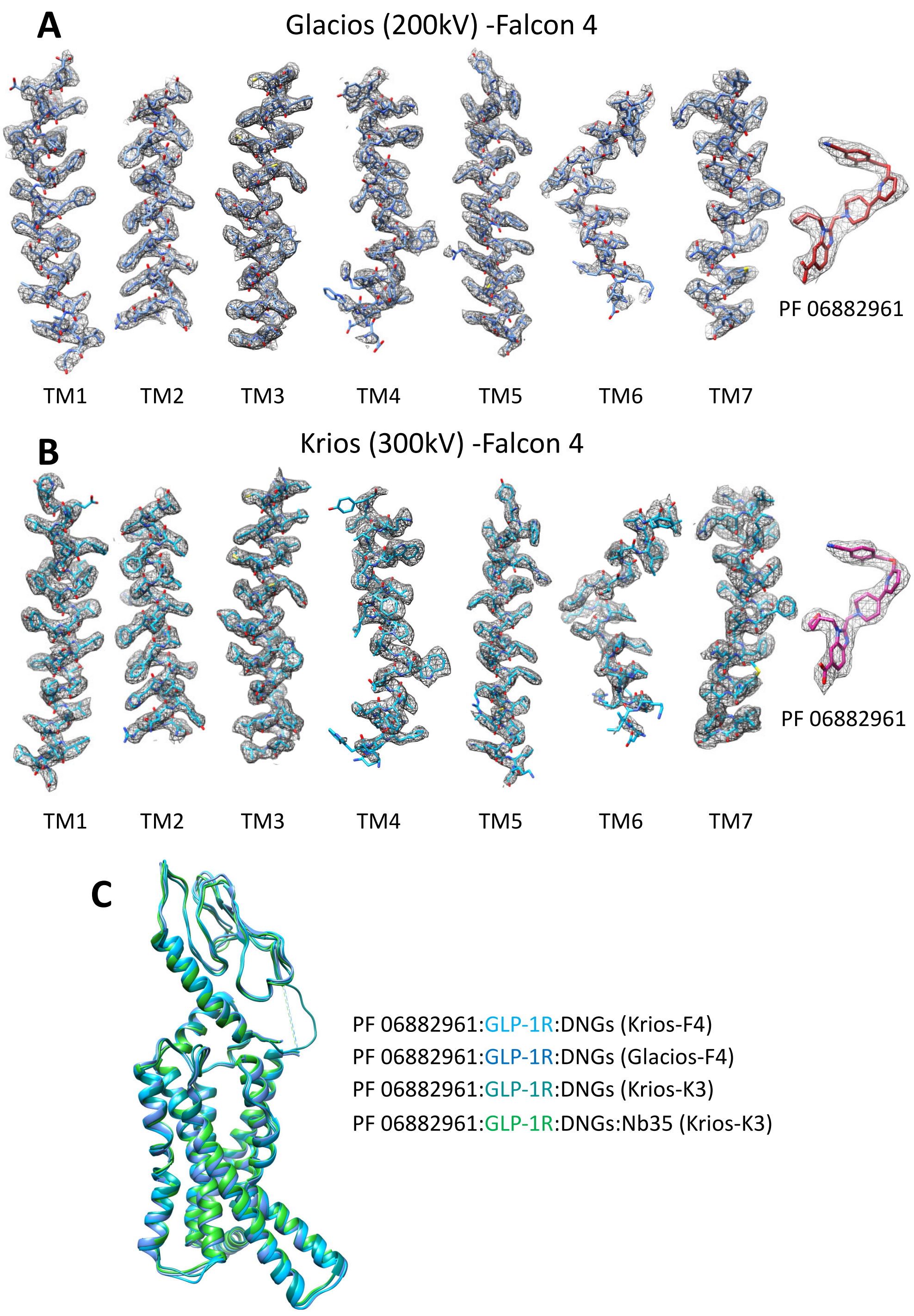


**Supplementary Figure 10. Representative atomic modelling into maps of the PF 06882961-GLP-1R-Gs complex imaged with 300kV (Krios-Falcon 4) or 200kV (Glacios-Falcon 4) cryo-EM.** (**A**) Data set from the 200kV Glacios and Falcon 4 system. (**B**) Data set from 300kV Krios and Falcon 4 system. Models of all seven TM helices the GLP-1R (blue, Glacios data; sky blue Krios data), and the ligand, PF 06882961 (dark red, Glacios data; dark pink, Krios data), are illustrated in the density map (mesh) generated from the receptor-focused maps via zone of 2 Å and mask on each component using UCSF Chimera. All residues are displayed in stick format. (**C**) Overlay of receptor backbone in ribbon format of the PF 06882961-GLP-1R structures determined in the absence of Nb35 (blue, Glacios-F4; sky blue, Krios-F4; cyan, Krios-K3) and in the presence of Nb35 (PDB 6X1A; green, Krios-K3).

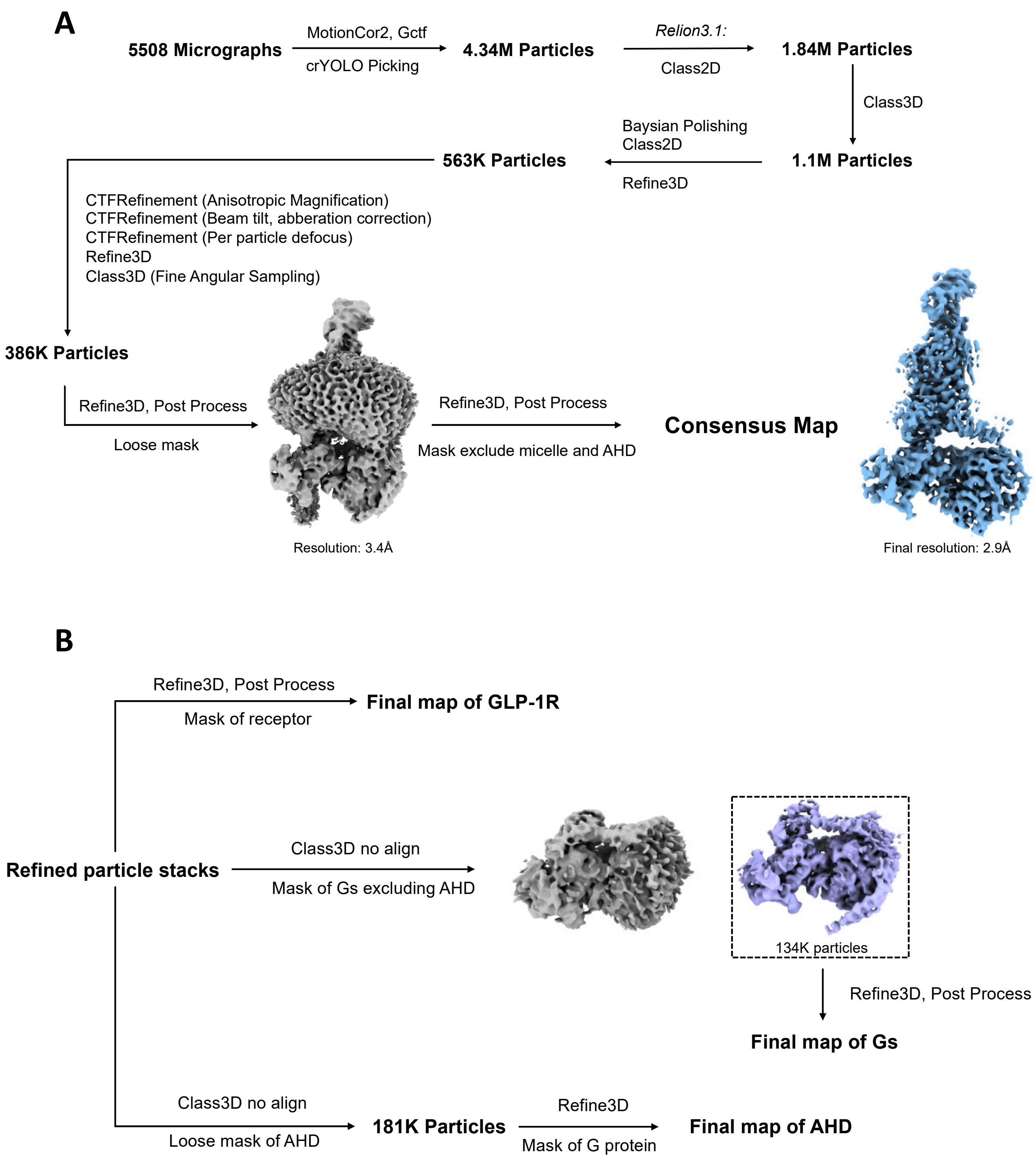


**Supplementary Figure 11. Cryo-EM data processing workflow for the main data set (300kV Krios – K3 detector).** (**A**) Reconstruction of the PF 06882961-GLP-1R-Gs consensus map. The 3D map of the whole particle is displayed in grey, and 3D map of the complex, masking out the micelle and AHD, is displayed in blue; (**B**) Local focused refinement, for the receptor, G protein and AHD. In the Gs-focused 3D classification, a representative class with Gγ density absent is displayed in grey and the best resolved class showing each subunit is displayed in purple.

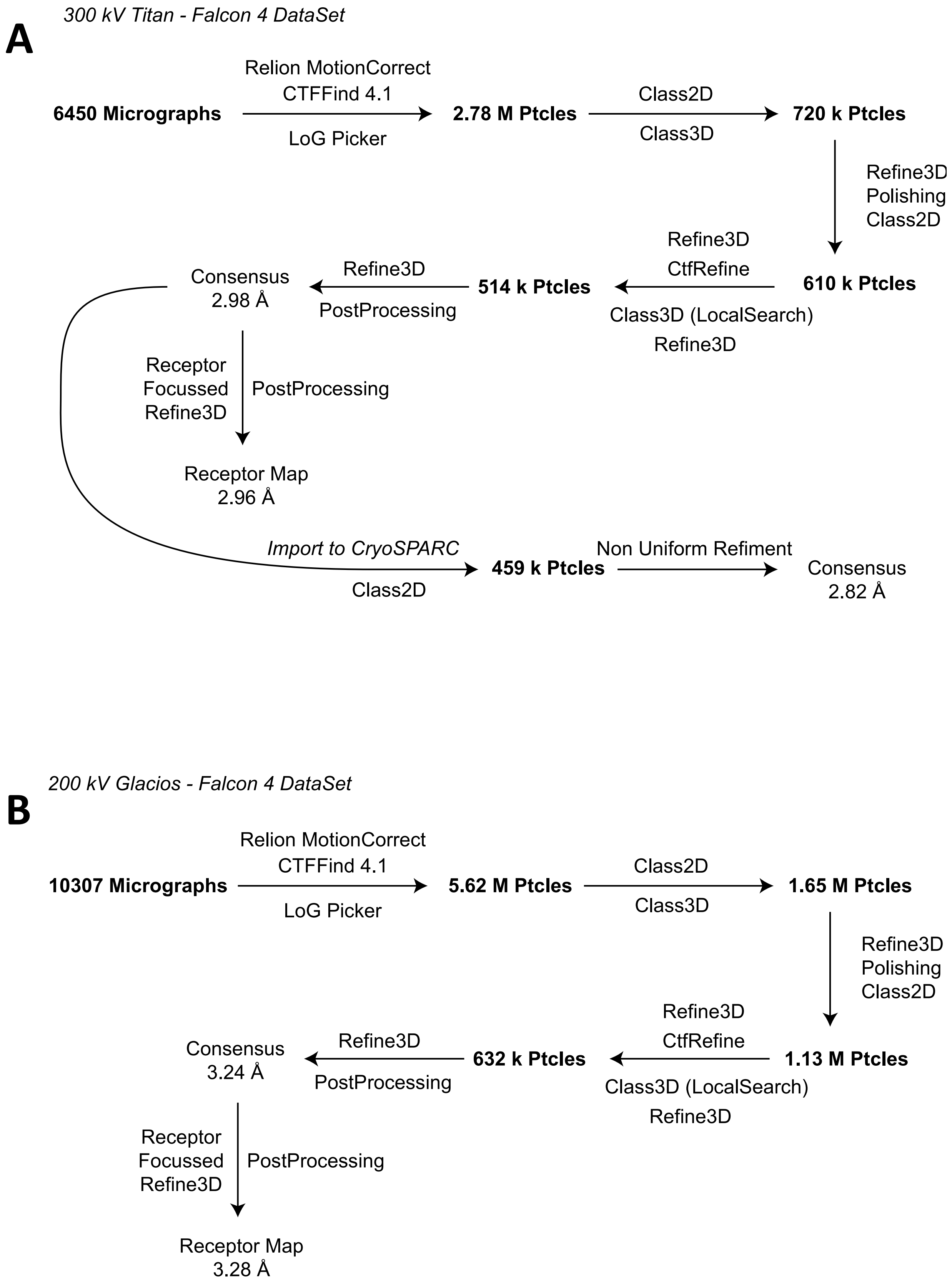


**Supplementary Figure 12. Cryo-EM data processing workflows for data collected on (A) 300kV (Krios-Falcon 4) or (B) 200kV (Glacios-Falcon 4) systems.**

**Supplementary Video Legends**

**Video S1. 3D variability analysis of PF 06882961-bound GLP-1R-Gs complexes.** CryoSPARC 3D variability analysis performed on the PF 06882961-GLP-1R-Gs complex in the presence or absence of Nb35. The top three principle motions are recorded. Transition 1: Principal component 1; Transition 2: Principal component 2; Transition 3; Principal component 3.

**Video S2. 3D variability analysis reveals differences in the conformational dynamics of the Gαs AHD versus in the absence versus presence of Nb35.** CryoSPARC 3D variability analysis of principal component 3 for both complexes shown at low contour. These illustrate the differences in the dynamics of the Gα AHD.

**Video S3. Morphs of the Gαs AHD between the inactive, GDP-bound, and GLP-1R-coupled conformations.** An overlay of inactive GDP-bound Gs (6EG8, ras-like domain in grey) and PF 06882961-bound complex (GLP-1R in blue and ras-like domain in gold) is displayed in ribbon format. A morph of the AHD (orange) between the two conformations relative to the ras-like domain is recorded. Proteins are displayed in ribbon format. GDP and Mg2+ (green) are displayed as spheres coloured by heteroatom.

**Supplementary Tables**

### Table S1: Interactions between the GLP-1R and PF 06882961 in the presence or absence of Nb35

| PF 06882961 functional groups | PF:GLP-1R:DNGs | PF:GLP-1R:DNGs:Nb35 | Interactions |
|---|---|---|---|
| Cyano-2-fluorobenzyl-oxy | L32$^{ECD}$<br>W33$^{ECD}$<br>V36$^{ECD}$<br>S206$^{ECL1}$<br>T207$^{ECL1}$<br>L217$^{ECL1}$<br>Q221$^{ECL1}$ | L32$^{ECD}$<br>W33$^{ECD}$<br>V36$^{ECD}$<br>S206$^{ECL1}$<br>T207$^{ECL1}$<br>L217$^{ECL1}$<br>Q221$^{ECL1}$ | Hydrophobic interaction<br>π-π interaction |
| Pyridine | S31$^{ECD}$<br>L141$^{1.36}$<br>F381$^{7.36}$ | S31$^{ECD}$<br>L141$^{1.36}$ | Hydrophobic interaction |
| Piperidine | L201$^{2.71}$<br>W203$^{2.73}$<br>F381$^{7.36}$ | L201$^{2.71}$<br>W203$^{2.73}$<br>F381$^{7.36}$ | Hydrophobic interaction<br>π-π interaction |
| Benzo-imidazole | K197$^{2.67}$ | K197$^{2.67}$ | H bond |
| | F230$^{3.33}$<br>M233$^{3.36}$<br>F381$^{7.36}$<br>L384$^{7.39}$ | F230$^{3.33}$<br>M233$^{3.36}$<br>F381$^{7.36}$<br>L384$^{7.39}$ | Hydrophobic interaction<br>π-π interaction |
| Oxetane | Q221$^{ECL1}$<br>F230$^{3.33}$<br>C296$^{ECL2}$<br>T298$^{ECL2}$ | Q221$^{ECL1}$<br>F230$^{3.33}$<br>C296$^{ECL2}$<br>T298$^{ECL2}$ | Hydrophobic interaction |
| Carboxylic acid | F230$^{3.33}$<br>L384$^{7.39}$ | F230$^{3.33}$ | Hydrophobic interactions |
| | R380$^{ECL3/7.35}$ | *R380$^{ECL3/7.35}$ | H bond |
| | Y152$^{1.47}$<br>R190$^{2.60}$<br>Q234$^{3.37}$<br>R299$^{ECL2}$ | Y152$^{1.47}$<br>R190$^{2.60}$<br>Q234$^{3.37}$<br>R299$^{ECL2}$ | Water mediated |

Interactions in their PDB files between PF 06882961 and the receptor were determined using Ligplot$^{+}$ (Laskowski and Swindells 2011). Hydrogen (H) bonds were additionally determined using UCSF Chimera. Residues labelled with * are hydrogen bond interactions not shown in Ligplot$^{+}$.

**Table S2: Interactions formed between GLP-1R and heterotrimeric Gs proteins**

| G protein subunit | G protein residue No. | GLP-1R PF:DNGs | GLP-1R PF:DNGs:Nb35 | GLP-1R GLP-1:DNGs:Nb35 |
|---|---|---|---|---|
| Gαs Ras αN | Q35$^{\alpha N}$ | | | S261$^{ICL2}$ |
| | A39$^{\alpha N}$ | | | V259$^{ICL2}$ |
| | H41$^{\alpha N}$ | F257$^{ICL2/3.60}$ | | F257$^{ICL2/3.60}$ |
| Gαs Ras β3 | V217$^{\beta 3}$ | F257$^{ICL2/3.60}$ | | |
| Gαs Ras α5 | F376$^{\alpha 5H}$ | F257$^{ICL2/3.60}$ | | F257$^{ICL2/3.60}$ |
| | C379$^{\alpha 5H}$ | | | F257$^{ICL2/3.60}$ |
| | R380$^{\alpha 5H}$ | F257$^{ICL2/3.60}$ | F257$^{ICL2/3.60}$ | F257$^{ICL2/3.60}$ |
| | D381$^{\alpha 5H}$ | K334$^{5.64}$ | K334$^{5.64}$ | K334$^{5.64}$ (H bond) |
| | I383$^{\alpha 5H}$ | | F257$^{ICL2/3.60}$ | F257$^{ICL2/3.60}$ |
| | Q384$^{\alpha 5H}$ | L255$^{3.58}$ (bb, H bond)<br>K334$^{5.64}$ (H bond) | L255$^{3.58}$ (bb, H bond)<br>K334$^{5.64}$ (H bond) | L255$^{3.58}$ (bb, H bond)<br>K334$^{5.64}$ (H bond) |
| | R385$^{\alpha 5H}$ | K334$^{5.64}$ (bb, H bond)<br>A337$^{ICL3/5.67}$<br>N338$^{ICL3}$ (H bond) | K334$^{5.64}$ (bb, H bond) | K334$^{5.64}$ (bb, H bond)<br>A337$^{ICL3/5.67}$<br>*N338$^{ICL3}$ |
| | H387$^{\alpha 5H}$ | L254$^{3.57}$ | L254$^{3.57}$ (bb, H bond) | L254$^{3.57}$ (Water mediated) |
| | L388$^{\alpha 5H}$ | L255$^{3.58}$<br>K334$^{5.64}$ | | L255$^{3.58}$<br>I330$^{5.60}$<br>*V331$^{5.61}$ |
| | Q390$^{\alpha 5H}$ | R176$^{2.46}$ | R176$^{2.46}$ | R176$^{2.46}$ |
| | Y391$^{\alpha 5H}$ | R176$^{2.46}$<br>E247$^{3.50}$ (Water mediated)<br>Y250$^{3.53}$ (π-π interaction)<br>L254$^{3.57}$ | E247$^{3.50}$ (Water mediated)<br>Y250$^{3.53}$ (π-π interaction)<br>L251$^{3.54}$ | R176$^{2.46}$<br>E247$^{3.50}$ (Water mediated)<br>Y250$^{3.53}$ (π-π interaction)<br>L251$^{3.54}$<br>T355$^{6.44}$ (Water mediated) |
| | E392$^{\alpha 5H}$ | R348$^{6.37}$ (H bond)<br>N406$^{8.47}$ | *N406$^{8.47}$ (H bond)<br>N407$^{8.48}$ (bb, H bond) | R348$^{6.37}$ (H bond)<br>N406$^{8.47}$ (H bond)<br>N407$^{8.48}$ |
| | L393$^{\alpha 5H}$ | R348$^{6.37}$<br>S352$^{6.41}$ | V331$^{5.61}$<br>S352$^{6.41}$ (H bond)<br>L356$^{6.45}$ | R348$^{6.37}$ (H bond)<br>S352$^{6.41}$ |
| | L394$^{\alpha 5H}$ | L335$^{5.65}$<br>R348$^{6.37}$ | K334$^{5.64}$<br>L335$^{5.65}$<br>R348$^{6.37}$ | K334$^{5.64}$<br>R348$^{6.37}$ |
| Gβ | R52$^{b}$ | R170$^{ICL1}$ (π-π interaction) | R170$^{ICL1}$ (π-π interaction) | R170$^{ICL1}$ (π-π interaction) |
| | V307$^{b}$ | | | L422$^{8.63}$ |
| | A309$^{b}$ | | R419$^{8.60}$ | R419$^{8.60}$ |

| | | |
|---|---|---|
| | | L422$^{8.63}$ |
| G310$^{b}$ | R419$^{8.60}$ | R419$^{8.60}$ |
| H311$^{b}$ | R419$^{8.60}$ | R419$^{8.60}$ |
| D312$^{b}$ | H171$^{ICL1}$<br>K415$^{8.56}$ (H bond) | *K415$^{8.56}$ (H bond)<br>*R419$^{8.60}$ (H bond) |

Interactions in the PDB files between Gs subunits and the receptor were determined using Ligplot$^{+}$. H bonds were additionally determined using UCSF Chimera. PF stands for PF 06882961. Subscript $^{\alpha 5H}$ indicates residues within α5 helix of Gαs; $^{\alpha N}$ indicates residues within αN helix of Gαs; $^{b}$ indicates residues within Gβ. Residues labelled with * are hydrogen bond interactions not shown in Ligplot$^{+}$.

**Table S3: Interactions formed between Gαs and Gβ proteins**

| Gαs regions | Gαs protein residue No. | Gβ of GLP-1R PF:DNGs | Gβ of GLP-1R PF:DNGs:Nb35 | Gβ of GLP-1R GLP-1:DNGs:Nb35 |
|---|---|---|---|---|
| αN | E16$^{\alpha N}$ | T86$^{b}$<br>N88$^{b}$ (H bond) | | T86$^{b}$ |
| | Q19$^{\alpha N}$ | D83$^{b}$ (H bond)<br>T86$^{b}$ (H bond)<br>N88$^{b}$ (H bond) | *D83$^{b}$ (H bond)<br>T86$^{b}$ (H bond)<br>N88$^{b}$ | D83$^{b}$ (H bond)<br>T86$^{b}$ (H bond)<br>N88$^{b}$ (H bond) |
| | R20$^{\alpha N}$ | | | T86$^{b}$<br>N88$^{b}$ |
| | N23$^{\alpha N}$ | K89$^{b}$ (bb, H bond) | N88$^{b}$<br>K89$^{b}$ (bb, H bond) | N88$^{b}$<br>K89$^{b}$ (bb, H bond) |
| | I26$^{\alpha N}$ | K89$^{b}$<br>A92$^{b}$ | K89$^{b}$ | K89$^{b}$<br>V90$^{b}$<br>H91$^{b}$<br>A92$^{b}$ |
| | E27$^{\alpha N}$ | | K89$^{b}$ (H bond) | K89$^{b}$ (H bond) |
| | L30$^{\alpha N}$ (no sc density) | G53$^{b}$<br>L55$^{b}$<br>I80$^{b}$ | G53$^{b}$<br>K78$^{b}$<br>K89$^{b}$ | G53$^{b}$<br>K78$^{b}$<br>K89$^{b}$ |
| | D33$^{\alpha N}$ | L55$^{b}$ | K78$^{b}$ | K78$^{b}$ (H bond) |
| | K34$^{\alpha N}$ | L55$^{b}$ | | L55$^{b}$ |
| | Y37$^{\alpha N}$ | A56$^{b}$ | L55$^{b}$<br>A56$^{b}$ | L55$^{b}$<br>A56$^{b}$ |
| | R38$^{\alpha N}$ | | | L55$^{b}$ (bb, H bond) |
| β2 | G206$^{\beta 2}$ | L117$^{b}$<br>N119$^{b}$ | L117$^{b}$<br>N119$^{b}$ | L117$^{b}$<br>N119$^{b}$ |
| | I207$^{\beta 2}$ | W99$^{b}$<br>L117$^{b}$<br>D118$^{b}$ | W99$^{b}$<br>D118$^{b}$ | W99$^{b}$ |
| β2/β3-loop | F222$^{\beta 2/\beta 3_loop}$ | W99$^{b}$ | W99$^{b}$ | W99$^{b}$ |
| β3/α2-loop | A226$^{\beta 3/\alpha 2_loop}$ | No sc desity | N119$^{b}$ (H bond) | T143$^{b}$ |
| | Q227$^{\beta 3/\alpha 2_loop}$ | No sc density | L117$^{b}$ (bb, H bond)<br>N119$^{b}$ (H bond)<br>Y145$^{b}$ (bb, H bond) | N119$^{b}$<br>G144$^{b}$<br>Y145$^{b}$ (bb, H bond) |
| | R228$^{\beta 3/\alpha 2_loop}$ | No sc density | G162$^{b}$ (bb, H bond)<br>D163$^{b}$<br>T164$^{b}$<br>D186$^{b}$ (H bond) | G162$^{b}$ (bb, H bond)<br>T164$^{b}$<br>D186$^{b}$ (H bond) |
| | R232$^{\beta 3/\alpha 2_loop}$ | C204$^{b}$ | C204$^{b}$<br>D228$^{b}$ (H bond) | C204$^{b}$<br>D228$^{b}$ (H bond) |
| | K233$^{\beta 3/\alpha 2_loop}$ | M101$^{b}$<br>Y145$^{b}$<br>M188$^{b}$<br>D228$^{b}$<br>N230$^{b}$ (H bond) | M188$^{b}$<br>C204$^{b}$<br>D228$^{b}$<br>N230$^{b}$ (H bond) | Y145$^{b}$<br>M188$^{b}$<br>C204$^{b}$<br>D228$^{b}$<br>N230$^{b}$<br>D246$^{b}$ (H bond) |

| | | | | |
|---|---|---|---|---|
| | W234$^{\beta3/\alpha2_loop}$ | L117$^{b}$ | L117$^{b}$ | L117$^{b}$ |
| α2 | Q236$^{\alpha2}$ | | | K57$^{b}$<br>Y59$^{b}$<br>R314$^{b}$ (H bond)<br>W332$^{b}$ |
| | C237$^{\alpha2}$ | *Y59$^{b}$ (H bond)<br>Q75$^{b}$<br>W99$^{b}$ | K57$^{b}$ (H bond)<br>W99$^{b}$ | K57$^{b}$ (H bond)<br>Y59$^{b}$<br>Q75$^{b}$<br>W99$^{b}$<br>M101$^{b}$ |
| | F238$^{\alpha2}$ | W99$^{b}$<br>L117$^{b}$ | W99$^{b}$<br>L117$^{b}$ | W99$^{b}$<br>L117$^{b}$ |
| | N239$^{\alpha2}$ | K57$^{b}$ (H bond)<br>W332$^{b}$ | K57$^{b}$ (H bond)<br>W332$^{b}$ | K57$^{b}$ (H bond)<br>W332$^{b}$ |
| | D240$^{\alpha2}$ | | | K57$^{b}$ (H bond) |
| α3/β5 -loop | K280$^{\alpha3/\beta5\ _loop}$ | | | D290$^{b}$ |
| | W281$^{\alpha3/\beta5\ _loop}$ | R314$^{b}$<br>W332$^{b}$ | D290$^{b}$<br>R314$^{b}$<br>W332$^{b}$ | D290$^{b}$<br>R314$^{b}$<br>W332$^{b}$ |

Interactions in the PDB files between Gαs and Gβ subunits were determined using Ligplot$^{+}$. H bonds were additionally determined using UCSF Chimera. Subscript $^{\beta2}$, $^{\beta2/\beta3\_loop}$, $^{\alpha2}$ or $^{\alpha3/\beta5\ \_loop}$ indicates residues within the corresponding region of Gαs subunit. Residues labelled with * are hydrogen bond interactions not shown in Ligplot$^{+}$.

**Table S4: Interactions formed between Nb35 and β3/α2 loop of Gαs subunit**

| **β3-loop-α2 residue No.** | **Nb35 residue No.** |
|---|---|
| $R228^{\beta3/\alpha2\_loop}$ | $T114^{n}$ (H bond) |
| $D229^{\beta3/\alpha2\_loop}$ | $S112^{n}$ |
| | $T113^{n}$ (H bond) |
| $E230^{\beta3/\alpha2\_loop}$ | $D109^{n}$ |
| | $S112^{n}$ |
| | $T114^{n}$ |
| | $Y115^{n}$ |
| $R231^{\beta3/\alpha2\_loop}$ | $D109^{n}$ (bb, H bond) |
| $R232^{\beta3/\alpha2\_loop}$ | $D109^{n}$ (H bond) |
| | $P100^{n}$ |
| | $Y115^{n}$ |
| $I235^{\beta3/\alpha2\_loop}$ | $F108^{n}$ |

Interactions in the PDB files between Nb35 and the β3/α2 loop were determined using Ligplot$^{+}$. Subscript $^{n}$ indicates residues within Nb35.

**Table S5: Interactions formed between Gβ and Gγ subunits**

| Gγ residue No. | Gβ of GLP-1R PF:DNGs | Gβ of GLP-1R PF:DNGs:Nb35 | Gβ of GLP-1R GLP-1:DNGs:Nb35 |
|---|---|---|---|
| S8$^{g}$ | L4$^{b}$ | L4$^{b}$ | L4$^{b}$ |
| I9$^{g}$ | E3$^{b}$<br>L4$^{b}$<br>L7$^{b}$ | E3$^{b}$<br>L4$^{b}$ | L4$^{b}$ |
| A12$^{g}$ | L7$^{b}$ | L7$^{b}$ | |
| R13$^{g}$ | L7$^{b}$ | L7$^{b}$ | L7$^{b}$ |
| V16$^{g}$ | L14$^{b}$ | A11$^{b}$<br>L14$^{b}$ | E10$^{b}$<br>L14$^{b}$ |
| Q18$^{g}$ | M217$^{b}$<br>C218$^{b}$ (bb, H bond) | C218$^{b}$ | C218$^{b}$ (H bond) |
| L19$^{g}$ | A11$^{b}$<br>L14$^{b}$<br>K15$^{b}$<br>I18$^{b}$ | A11$^{b}$<br>L14$^{b}$ | A11$^{b}$<br>L14$^{b}$<br>I18$^{b}$ |
| K20$^{g}$ | L14$^{b}$ | L14$^{b}$ | L14$^{b}$ |
| M21$^{g}$ | | | M217$^{b}$ |
| E22$^{g}$ | I18$^{b}$<br>Q220$^{b}$<br>T221$^{b}$ (bb, H bond) | I18$^{b}$<br>C218$^{b}$<br>R219$^{b}$<br>Q220$^{b}$<br>T221$^{b}$ | C218$^{b}$<br>R219$^{b}$<br>Q220$^{b}$<br>T221$^{b}$ (H bond) |
| A23$^{g}$ | Q17$^{b}$<br>I18$^{b}$ | I18$^{b}$ | Q17$^{b}$<br>I18$^{b}$ |
| I25$^{g}$ | D258$^{b}$ | D258$^{b}$ | D258$^{b}$ |
| R27$^{g}$ | A21$^{b}$<br>*C25$^{b}$ (H bond)<br>R256$^{b}$<br>D258$^{b}$ (H bond) | I18$^{b}$<br>A21$^{b}$<br>R22$^{b}$<br>C25$^{b}$<br>R256$^{b}$ | A21$^{b}$<br>C25$^{b}$<br>R256$^{b}$<br>D258$^{b}$ (H bond) |
| I28$^{g}$ | C25$^{b}$<br>R256$^{b}$ | C25$^{b}$ (H bond)<br>R256$^{b}$<br>A257$^{b}$ | C25$^{b}$<br>R256$^{b}$ (bb, H bond)<br>A257$^{b}$ |
| K29$^{g}$ | C25$^{b}$ | C25$^{b}$<br>D27$^{b}$ | A24$^{b}$ (bb, H bond)<br>C25$^{b}$<br>D27$^{b}$ |
| V30$^{g}$ | C25$^{b}$ (bb, H bond)<br>A26$^{b}$<br>D27$^{b}$<br>A28$^{b}$<br>Q259$^{b}$<br>L261$^{b}$ | C25$^{b}$ (bb, H bond)<br>A26$^{b}$<br>D27$^{b}$<br>A28$^{b}$<br>Q259$^{b}$ | C25$^{b}$ (bb, H bond)<br>A26$^{b}$<br>D27$^{b}$<br>A28$^{b}$<br>Q259$^{b}$<br>L261$^{b}$ |
| S31$^{g}$ | D27$^{b}$ (H bond) | D27$^{b}$ | D27$^{b}$ (H bond) |
| A33$^{g}$ | D254$^{b}$<br>A257$^{b}$ | D254$^{b}$<br>R256$^{b}$ | D254$^{b}$ |

| | | | |
|---|---|---|---|
| A34[g] | | L30[b]<br>I33[b]<br>L261[b] | L30[b] |
| A35[g] | | I33[b] | |
| D36[g] | | | R256[b] |
| L37[g] | F235[b]<br>L252[b]<br>L261[b] | F235[b]<br>L252[b]<br>L261[b] | F235[b]<br>N237[b]<br>L261[b] |
| M38[g] | L30[b]<br>I33[b]<br>T34[b]<br>L300[b] | L300[b] | I33[b]<br>T34[b] |
| Y40[g] | F235[b]<br>P236[b]<br>N237[b]<br>S281[b] | P236[b]<br>N237[b] | F235[b]<br>P236[b]<br>N237[b]<br>S281[b] |
| C41[g] | F235[b]<br>S281[b] (H bond)<br>L300[b] | R283[b]<br>L300[b] | F235[b]<br>S281[b]<br>G282[b]<br>R283[b]<br>L300[b] |
| E42[g] | | I37[b] | I37[b] |
| H44[g] | S281[b] | S281[b] | S281[b] |
| A45[g] | | S281[b] | |
| E47[g] | K280[b] (H bond) | | K280[b] |
| D48[g] | S279[b] (H bond)<br>S281[b] (bb, H bond) | S279[b]<br>S281[b] | S279[b] (H bond)<br>S281[b] (H bond) |
| P49[g] | D323[b]<br>G324[b]<br>M325[b] | G324[b]<br>M325[b] | D323[b]<br>G324[b]<br>M325[b] |
| L50[g] | I43[b]<br>G324[b]<br>M325[b]<br>V327[b]<br>N340[b] | I43[b]<br>M45[b]<br>G324[b]<br>M325[b]<br>V327[b] | I43[b]<br>L284[b]<br>G324[b]<br>M325[b]<br>V327[b] |
| L51[g] | V40[b]<br>R283[b] | I43[b]<br>R283[b] | V40[b]<br>R283[b]<br>L284[b] |
| V54[g] | | | M325[b] |
| E58[g] | | | M325[b] |
| N59[g] | N340[b] | N340[b] | N340[b] |
| P60[g] | R49[b]<br>Y85[b] | R49[b] (H bond)<br>Y85[b] | Y85<br>M325 |
| F61[g] | R48[b] | R48[b] | R48[b] |

| | | | |
|---|---|---|---|
| | R49$^{b}$<br>S84$^{b}$<br>Y85$^{b}$<br>A326$^{b}$<br>I338$^{b}$<br>N340$^{b}$ | S84$^{b}$<br>Y85$^{b}$<br>A326$^{b}$<br>I338$^{b}$<br>N340$^{b}$ | R49$^{b}$<br>S84$^{b}$<br>Y85$^{b}$<br>A326$^{b}$<br>I338$^{b}$<br>N340$^{b}$ |
| R62$^{g}$ | R48$^{b}$ | | |

Interactions in the PDB files between Gαs and Gβ subunits were determined using Ligplot$^{+}$. H bonds were additionally determined using UCSF Chimera. Subscript $^{g}$ indicates residues within the Gγ subunit. Residues labelled with * are hydrogen bond interactions not shown in Ligplot$^{+}$.

**Table S6: Data collection and refinement statistics.**

| **Data Collection** | PF 06882961:GLP-1R:DNGs **300 kV – K3** | PF 06882961:GLP-1R:DNGs **300 kV – F4** | PF 06882961:GLP-1R:DNGs **200 kV – F4** |
|---|---|---|---|
| Micrographs | 5508 | 6450 | 10307 |
| Electron dose ($e^-/A^2$) | 61.8 | 60 | 55.8 |
| Voltage (kV) | 300 | 300 | 200 |
| Pixel size (Å) | 0.83 | 0.82 | 0.95 |
| Defocus range (µm) | 0.5-1.5 | 0.5-1.5 | 0.5-1.5 |
| Symmetry imposed | C1 | C1 | C1 |
| Particles (final map) | 386 k | 459 k | 632 k |
| Resolution (0.143 FSC)(Å) | 2.9 | 2.82 | 3.24 |
| **Refinement** | | | |
| $CC_{map\_model}$ | | | |
| Map sharpening B factor (Å$^2$) | -76 | -103 | -118 |
| **Model Quality** | | | |
| R.m.s. deviations | | | |
| Bond length (Å) | 0.004 | 0.006 | 0.009 |
| Bond angles (°) | 0.726 | 0.703 | 0.894 |
| Ramachandran | | | |
| Favoured (%) | 96.67 | 93.01 | 94.56 |
| Outliers (%) | 0 | 0 | 0 |
| Rotamer outliers | 0 | 0 | 0 |
| C-Beta deviations (%) | 0 | 0 | 0 |
| Clashscore | 10.38 | 6.08 | 5.15 |
| MolProbity score | 1.75 | 1.78 | 1.65 |
